# Leveraging the Red List of Ecosystems for national action on coral reefs through the Kunming-Montreal Global Biodiversity Framework

**DOI:** 10.1101/2024.02.18.580850

**Authors:** Mishal Gudka, David Obura, Eric Treml, Melita Samoilys, Swaleh Aboud, Kennedy Osuka, James Mbugua, Jelvas Mwaura, Juliet Karisa, Ewout Knoester, Peter Musila, Mohamed Omar, Emily Nicholson

**Affiliations:** University of Melbourne, Grattan Street, Parkville, Victoria, 3010, Australia; CORDIO East Africa, 9 Kibaki Flats, Kenyatta Beach, Bamburi, Mombasa, Kenya; Pwani University, School of Pure & Applied Sciences, Kilifi, Kenya; Australian Institute of Marine Science (AIMS) and UWA Oceans Institute, The University of Western Australia, MO96, 35 Stirling Highway, Crawley, WA 6009; Department of Earth, Ocean and Ecological Sciences, University of Liverpool, Liverpool, England; Kenya Marine and Fisheries Research Institute, Mombasa, Kenya; Marine Animal Ecology, Wageningen University & Research, Wageningen, The Netherlands; A Rocha Kenya, Watamu, Kenya; Wildlife Research Training Institute, Naivasha, Kenya

**Keywords:** ecosystem assessment, global policy, ecological indicators, ecosystem management, decision support, biodiversity

## Abstract

Countries have committed to conserving and restoring ecosystems after signing the Kunming-Montreal Global Biodiversity Framework (GBF). The IUCN Red List of Ecosystems (RLE) will serve as a headline indicator to track progress of countries towards achieving this goal, and to guide action across the GBF’s targets. Using Kenyan coral reefs, we demonstrate how nations implementing the GBF, can use standardised estimates of ecosystem degradation from RLE assessments to support site-specific management decisions. We undertook a reef-by-reef analysis to evaluate the relative severity of decline of four key ecosystem components over the past 50-years: hard corals, macroalgae, parrotfish and groupers. Using the two benthic indicators, we also calculated standardised estimates of state to identify reef sites which maintain a better condition through time relative to adjacent sites. Over the past 50 years, Kenya’s coral reefs have degraded across all four ecosystem components. At more than half the monitored sites both parrotfish and grouper abundance declined by more than 50%, while coral cover and macroalgae-coral ratio declined by at least 30%. This resulted in a Vulnerable threat status for coral reefs in Kenya based on degradation (under criterion D of the RLE). The temporal trends in coral cover revealed four sites which maintained an above average condition over their monitoring history (15-25 years). The results can guide management actions to contribute to at least nine of the 23 GBF targets. For example, we identified several sites with relatively healthy benthic and fish communities as candidate areas for protection measures under Target 3. We also found that Marine Protected Areas and Locally Managed Marine Areas, which restrict fishing and control gears, had lower declines in groupers compared to unmanaged areas, providing further evidence for their expansion. The RLE has a key role to play in monitoring and meeting the goals and targets of the Global Biodiversity Framework, and our work demonstrates how using the wealth of data within these assessments can inform local-scale ecosystem management and amplify the GBF’s impact.

## Introduction

The need to address ongoing loss of ecosystems has been recognised through an explicit goal to conserve ecosystems in the Kunming-Montreal Global Biodiversity Framework (GBF) of the UN Convention on Biological Diversity (CBD, 2022). This multi-lateral treaty on biodiversity comprises four goals and 23 targets which identify actions countries must undertake to achieve the goals, as well as a monitoring framework to track progress. While global ecosystem-based policy frameworks set aspirations, it is ultimately up to countries and local communities to make policies, decisions, and take action to achieve those goals (Obura et al., 2023). These decisions should be informed by the GBF but there are barriers which could inhibit this. For example, robust and informative ways for downscaling global datasets for national use are currently lacking (Buschke et al., 2023). Additionally, specific linkages between the GBF and desired national and sub-national policy and decision-making processes are underdeveloped (Carroll et al., 2022; Nicholson et al., 2024).

The International Union for Conservation of Nature (IUCN) Red List of Ecosystems (RLE) is the global standard for measuring the relative risk of collapse of ecosystem types (Keith et al., 2013). RLE assigns ecosystem types to risk categories (e.g., Endangered, Vulnerable) based on criteria relating to change in area and degradation. The RLE serves multiple roles in the monitoring framework of the GBF. This includes as a headline indicator to track progress of countries towards achieving Goal A (Nicholson et al., 2024), on enhancing ecosystem integrity, connectivity and resilience, and increasing ecosystem area (Joly, 2023). It is also a headline or complementary indicator for multiple targets (Nicholson et al., 2024). Thus, countries should undertake RLE assessments of their ecosystems for national reporting to the GBF. This provides an important opportunity to make the wealth of ecosystem data and information collected for national reporting useful for ecosystem management decisions at local levels (Nicholson et al., 2024). Previously, ecosystem-specific RLE assessments have been used for local scale ecosystem management, such as zonation of Burullus Protected Area in Egypt for a Biosphere Reserve nomination (Ghoraba et al., 2021), and fencing of Fathala forest, Senegal to increase protection (Keith et al., 2023). Ecosystem management and conservation can be scaled up through the GBF by developing priorities and action plans that contribute to meeting GBF targets on protection, sustainable use and restoration.

In the regional RLE assessment of the Western Indian Ocean (WIO) coral reefs, Kenyan coral reefs were assessed as part of the Northern Tanzania–Kenya ecoregion. This ecoregion was found to be Vulnerable due to biotic degradation (Obura et al., 2021). The same principles, data and indicators of key biotic processes and interactions (related to coral cover, macroalgal cover and abundance of parrotfish and groupers) can be used at the reef scale and finer in Kenya. This is the spatial scale where in-situ data collection happens, and where management actions are usually applied (McClanahan et al., 2005; Ban et al., 2011; Obura, 2013; Maina et al., 2015). We evaluate the relative severity of biotic degradation at each reef site over the past 50 years, and assess the national collapse risk for coral reefs under Criterion D. In addition, we conduct a complementary analysis from time-series data to identify and classify reef sites based on their potential resistance and recovery capacities to disturbances. We develop and test an efficient and transferable method to recommend potential management actions at the reef site scale linked to nine relevant GBF targets. In doing so, we showcase how all Parties to the CBD can effectively employ RLE assessments to inform and enhance local management decisions as they implement the GBF. To support application of our approach to other countries and ecosystems, we provide guidance on methods of aggregation, indicator selection and application of outputs to inform management.

## Methods

### Study area, data and indicators

Kenyan coral reefs extend almost continuously along the entire coastline of the country, with one break due to two major river flows: River Tana and Sabaki (McClanahan et al., 2000; Obura, 2001). The fringing reefs are present in four of the five coastal Counties (local government administrative units) (Fig. 1, Fig. S1), where they provide valuable coastal protection, and critical habitat for biodiversity that supports small-scale fishers and tourism activities (Cinner et al., 2009; Ransom and Mangi, 2010; Darling, 2014; Obura et al., 2017b). The reefs are impacted by fishing, extraction, pollution from various terrestrial sources and coral bleaching, with regulations and interventions implemented to try to manage these threats e.g., five national Marine Protected Areas (MPAs) and over twenty Locally Managed Marine Areas (LMMAs) (Kawaka et al., 2017; United Nations Environment Programme and Western Indian Ocean Marine Science Association, 2021).

**Figure 1:**
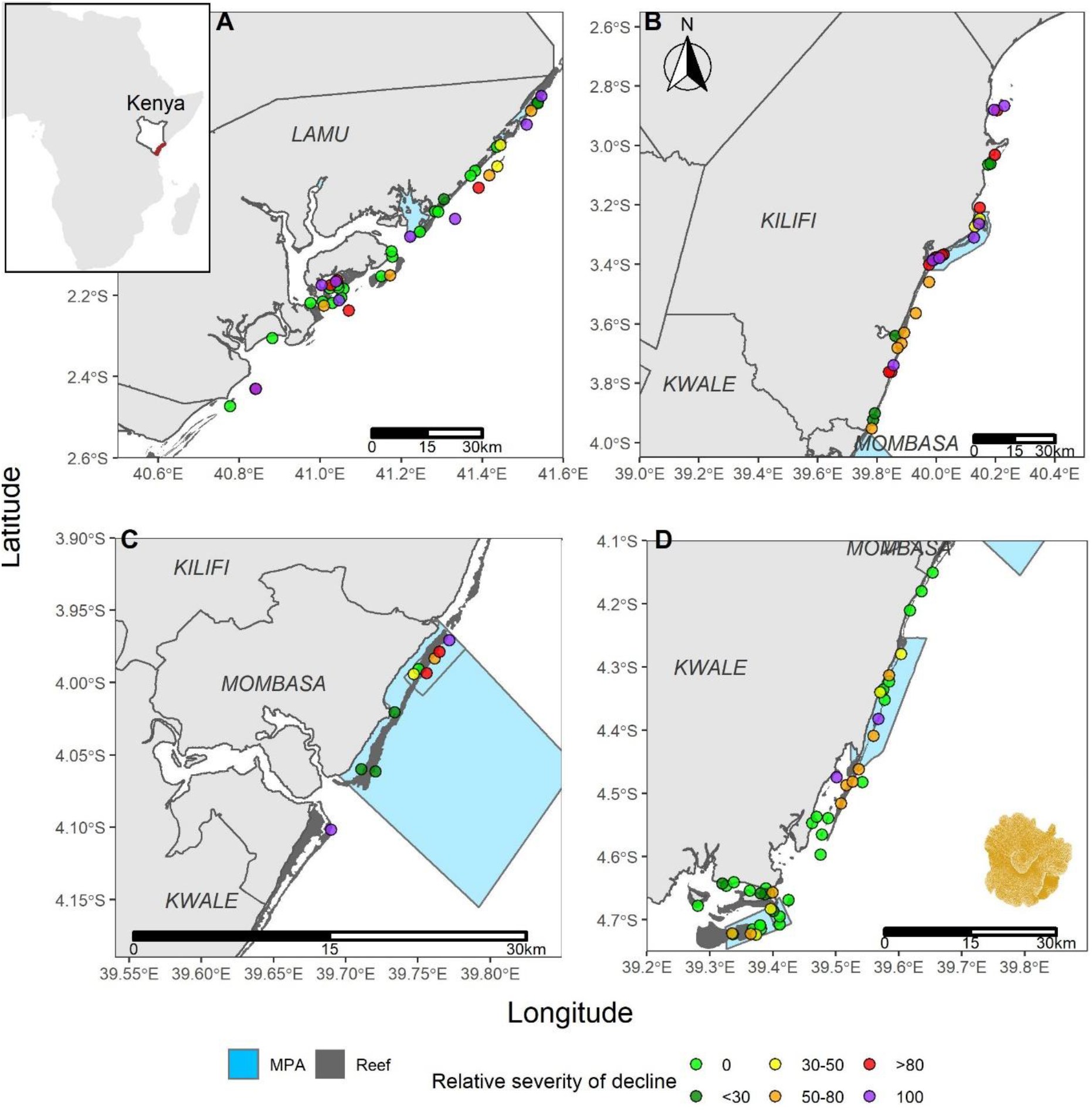
Relative severity of decline classifications over the past 50-years for hard coral cover at coral reef monitoring sites in the four coastal counties of Kenya (A) Lamu, B) Kilifi, C) Mombasa, D) Kwale – north to south). Light blue areas are national Marine Protected Areas (no-take zones and reserves), and dark grey areas represent coral reefs. Sites are coloured based on median relative severity of decline. Inset map on top-left corner shows Kenya with coastal counties highlighted in red.

Monitoring of coral reefs in Kenya and the WIO has employed underwater visual census methods to collect consistent data on benthic cover and fish abundance (Obura, 2013; Obura et al., 2017a). For this study we used a database consisting of long-term monitoring data (1987 - 2023), compiled by the WIO regional node of the Global Coral Reef Monitoring Network (GCRMN)(Souter et al., 2021). Data was analysed at the scale of a monitoring site, which is a fixed survey location within a reef zone or depth range covering an area of 10–250 m^2^.

Following RLE assessments for coral reefs in the WIO (Obura et al., 2021) and other regions (Bland et al., 2017; Uribe et al., 2021), as well as state and resilience assessments (Dahlgren et al., 2016; Cowburn et al., 2019; Gibbs and West, 2019), we assess four ecosystem components integral to the ecological functioning of coral reefs in Kenya: i) hard corals as the ecosystem engineers, creating structure and habitat, ii) macroalgae, as the principal competitor to coral, iii) herbivores which provide top-down control on algae, and iv) piscivores, which impose higher-order trophic effects. We selected the following indicators to represent these four components:

1. *Hard coral cover:* the percent cover of hard corals (%). Hard coral cover has historically been the primary measure of the general state (or health) of reefs.
2. *Macroalgae-coral ratio*: calculated as macroalgae cover/ (macroalgae + hard coral cover). The comparative levels of these two groups are important to gauge the level of competition between macroalgae and corals.
3. *Parrotfish abundance:* number of individuals of parrotfish (Scarinae within family Labridae) within a standard area (ind/ha). Parrotfish are a good proxy indicator for herbivory, as nearly all parrotfish species are grazers, and they comprise almost half of all herbivorous reef fish species in the WIO (Samoilys et al., 2019a).
4. *Grouper abundance:* number of groupers (family: Epinephelidae) within a standard area (ind/ha). Groupers are top predators on reefs and are also sensitive to fishing. Groupers represent over 80% of piscivore species in the WIO (Samoilys et al., 2019a).

### Relative severity of biotic degradation

In this study, we assess RLE Criterion D which focuses on the disruption of characteristic biotic processes and interactions of an ecosystem. This is assessed by quantifying how much a reef’s current state has changed from an initial state (e.g. 50 years ago) towards a threshold defining collapse (Keith et al., 2013; Murray et al., 2016). This is known as the ‘relative severity of decline’. By scaling indicators in this way, different types of indicators can be directly compared. We established initial and current values, and collapse thresholds for each indicator, and used them to calculate the relative severity of decline of reef sites (detailed description provided below)(Obura et al., 2021).

### Initial values

As per the RLE standard, initial values represent the indicator value 50 years ago (∼ 1970). Because monitoring data does not extend this far back, we followed the methods described in Obura et al., 2021 and Gudka et al., 2023, to estimate initial conditions (mean and standard deviation) for each indicator using data from select sites. Ideally, the initial value for an individual site should represent conditions in that area as closely as possible. Kenyan reefs were divided into five ecologically distinct zones consisting of between 24.5 – 269.2 km^2^ of reef area (Table S1, Fig. S1): Lamu-Kiunga, Malindi-Watamu, Mombasa-Kilifi, Diani-Chale and Msambweni-Kisite (Samoilys et al., 2017). Each zone represents the finest spatial resolution possible to enable robust estimates of initial values given the available historical data. For both benthic indicators, we used coral and macroalgae cover data from sites known to be in healthy condition prior to (or just after) the 1998 mass coral bleaching event, which marks a period prior to major, broad-scale disturbances in Kenya (n=74 for coral cover, n=66 for macroalgae-coral ratio). The exception was Diani-Chale, which was known to be degraded around 1970 (Fig. S1). We averaged site values in each zone to get sub-national zone means and ran a bootstrapping approach with replacement to generate standard deviations around the mean (Rowland et al., 2021; Gudka et al., 2023). Fishing pressure in 1970 was considerably lower compared to present day, so to approximate initial values corresponding to healthier fish populations, we included reference data from two remote and well protected sites in the Indian Ocean - Chagos Archipelago and Iles Glorieuses (Kaunda-Arara et al., 2003; Tuda and Wolff, 2015; Samoilys et al., 2017; Obura et al., 2021). For each sub-national zone, these values were combined with data from sites within MPAs (no-take zones or reserves – areas where regulated artisanal/traditional fishing is allowed) with at least six years of strong enforcement. We identified suitable MPA sites in each sub-national zone except Tiwi-Chale, where we applied the initial values from Mombasa-Kilifi as they have the most similar geomorphology and reef types (Andréfouët et al., 2009; Samoilys et al., 2019a). We excluded 43 data points (3% of total data) with exceptionally high values to address possible observer errors (mean >900 indiv/ha for groupers and >1500 indiv/ha for parrotfish, with a 10% buffer). These limits were informed by our reference sites and surveys elsewhere.

### Current values

We used a period spanning the last 11 years (2013-2023) to calculate current values because a) sites varied in the year of most recent data, b) to smooth out inter-annual inconsistencies and fluctuations (e.g., due to bleaching in 2016), and c) to increase the sample size of qualifying sites (Obura et al., 2021). We then averaged all points within this timeframe for sites with multiple data points.

### Collapse threshold

The collapse threshold is the level at which the measured process or feature stops functioning, causing severe and persistent degradation to the ecosystem (Bland et al., 2018; Rowland et al., 2018). We applied collapse thresholds used in the regional RLE (Obura et al., 2021) to make results comparable: 5% for hard coral cover; 0.833 for macroalgae-coral ratio; and 10% and 20% of the initial population values in each sub-national zone for parrotfish and grouper, respectively. For the fish indicators, similarities in the geomorphology of the reefs in sub-national zones of Malindi-Watamu, Kilifi-Mombasa and Tiwi-Chale suggest they should collapse at the same level despite having different initial values. Therefore, sites in these three zones were considered together to calculate a single collapse threshold for the entire area.

### Calculating relative severity of decline

For each indicator, we assessed the relative severity of decline for each reef site using the formula (Murray et al., 2016):

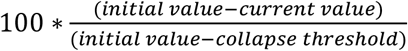

A value of 0 (or less) indicates a reef has maintained or improved condition, while a reef with a value of 100 (or greater) has collapsed. To capture the uncertainty in the initial value estimates (bootstrapped standard deviation), we calculated relative severity 750 times for each site, each time randomly sampling an initial value from the bootstrapped distribution (assuming a normal distribution). We rounded relative severity values to 100 and 0 to make results easier to interpret since this doesn’t affect the calculation of the risk categories (see next section). If we needed a single relative severity of decline value for each site, we used the median value across all iterations.

We undertook a comparative assessment of severity of decline for each indicator across existing area-based management regimes (reserves (partially restricted areas), open-access areas, and no-take zones) using a Kruskal-Wallis rank sum test.

### National risk status under Criterion D

The collapse risk status under Criterion D was determined in three steps. First, we assigned a risk status for each iteration using the proportion of sites (extent) in each relative severity bin (>30, >50, >80) (see extent vs relative severity matrix Fig. S2). Then, the risk status for each indicator is determined by summarising the percentage of iterations returning each risk level e.g., if 50% of iterations were Least Concerned, and 50% were Vulnerable, the overall status for that indicator can be assumed as Near Threatened. As per RLE guidelines, we considered the final risk status of the ecosystem as the most threatened status across all indicators. For example, if reefs are Least Concerned for corals, Near Threatened for macroalgae-coral ratio, Least Concerned for parrotfish and Vulnerable for groupers, then the overall risk status for that site would be Vulnerable based on groupers. We decided to limit the risk status of fish indicators to a maximum of Vulnerable to temper their effect on the final threat status because: i) we had lower confidence in the initial values and collapse thresholds (indicated by the large standard deviations for each sub-national zone – Table S2), ii) if parrotfish or grouper abundance is low, a reef’s functioning and resilience will be degraded, but the reef would not be ‘collapsed’, and iii) it emphasises the importance of the foundational, keystone taxa, hard corals.

Detailed analysis of reef site trajectory

We followed the methods in Guest et al., 2018 and analysed time-series data of the two benthic indicators to determine site state relative to all other sites, along with variability in state through time. We standardised estimates of each indicator as Z-scores, which quantifies how much, and in what direction (positive or negative), each site differs from a national mean value (Guest et al., 2018). Because sites were not surveyed each year, we used bins across multiple years to calculate national means: pre-1998, 1998 – 2003, 2004 – 2008, 2009 – 2015, 2016 – 2023. The bins were selected to split the data somewhat equally but also to combine periods of similar conditions, and account for the two major bleaching events in 1998 and 2016. We averaged multiple samples from a site within the bin, then averaged these values to calculate the national bin mean and standard deviation. All sites including sites with a single survey point were included in national bin calculations.

For each year a site is surveyed, a Z-score for that site is calculated by dividing the difference between the site mean for that year and the corresponding national bin mean, by the standard deviation of that bin (Guest et al., 2018):

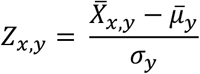

where Z_x,y_ is the Z-score for site x in year y, 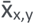 is the indicator mean for site x in year y, and 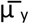 and σ_y_ are the national (binned) mean and standard deviation corresponding to year y respectively.

To meaningfully detect trends, a site was only assessed if it had a minimum of 4 years of data spread across at least 2 time-periods. This is consistent with regional status reports (Obura et al., 2017a) and allowed us to include more sites, while still providing enough samples for a robust analysis. In addition, a site needed to have data in the most recent time-period (2016-2023) to examine responses to the 2016 bleaching event. We inverted the scores calculated for macroalgae-coral ratio to ensure positive Z-scores correspond to healthier reef condition. The median Z-score for each site was calculated across all surveyed years (median is less sensitive to extreme values and fluctuations) (Guest et al., 2018).

We also examined the inter-annual variation for each site using the standard deviation of mean indicator values across a site’s time-series. Sites were classified as having low, moderate or high variance by dividing the range of standard deviation values across all sites into thirds.

We identified two outlier points which had abrupt and ecologically unrealistic inter-annual fluctuations in indicator values. Outliers exhibited a positive inter-annual rate of change exceeding 2.5 standard deviations from the average (calculated across all positive changes from all sites) (Dunn, 2021), and exceeded the 75th percentile of all other values for that site by 1.5 times the inter-quartile range. We winsorized outlier values, which involved bringing the value down to the next highest value (Rivest, 1994; Wu and Zuo, 2009). The analysis was completed using these corrected data. The results were visualised in a bi-plot of median Z-scores against standard deviation.

### Characterisation of site resilience

We defined five distinct classifications of possible trajectories of reefs that could be reliably interpreted from the analysis. We adapted phase-shifted, rebound/recovered and resistant from Guest et al. 2018 and introduced degraded and declined as two additional classifications. We formulated and assigned quantitative thresholds of variables from the analysis to each classification to represent the expected trajectory. The variables were the site: standard deviation, median Z-score, first Z-score, most recent Z-score, direction of the most recent Z-score (derived by subtracting the recent and median values), and maximum and minimum Z scores.

### Supporting local management decisions

We developed a two-dimensional phase-space to analytically link the benthic conditions and fish populations at sites to local management strategies related to GBF targets. We used a Principal Component Analysis (PCA) to reduce the four relative severity of decline indicators into two axes. This allows all four ecosystem components to be considered without masking their individual levels, as would be the case if combined into an index. The space is divided into 4 quadrants based on the median values of each PC. The framework helps to identify actions that shift sites located anywhere in the space toward improved benthic and/or fish states or maintain them in a ‘healthy’ condition. Management actions proposed were based on our knowledge of the local context of the reef sites and threats they face and were related to relevant GBF targets (T1–spatial planning, T2–restoration, T3– protection, T4– recovering threatened species, T5– wild species use, T7– Reduce pollution, T8–climate adaptation, T9– benefits and tourism).

## Results

### National threat status under Criterion D

Kenya’s reefs were classified as Vulnerable under Criterion D of the RLE. This was primarily driven by declines in grouper and parrotfish abundance over the past 50 years, which were both classified as Vulnerable. Both benthic indicators were assessed as Near Threatened, though macroalgae-coral ratio iterations returned this result only 45% of the time (51% as Least Concerned).

### Relative severity of biotic degradation

The relative severity of decline varied across sites (e.g. Fig. 1 for corals) and across the four indicators (hard coral, macroalgae-coral ratio, groupers and parrotfish). The biggest decline was in groupers, with a median relative severity of decline of 100% (n=105 sites) (Fig. 2), compared to 52% for parrotfish (n=102), 33% for hard corals and 43% for macroalgae-coral ratio (n=132 for hard coral and macroalgae). Fifty-one sites showed similar or improved condition compared to initial levels (relative severity of zero) for hard coral cover, 32 sites for macroalgae-coral ratio, 23 sites for parrotfish and 6 sites for groupers (Fig. 2). On the other end of the spectrum, a total of 19 sites were considered collapsed based on a hard coral cover less than 5%, and 16 sites based on macroalgae-coral ratio of 0.833. 19 sites were collapsed based on parrotfish abundance, and 64 sites based on grouper abundance.

**Figure 2:**
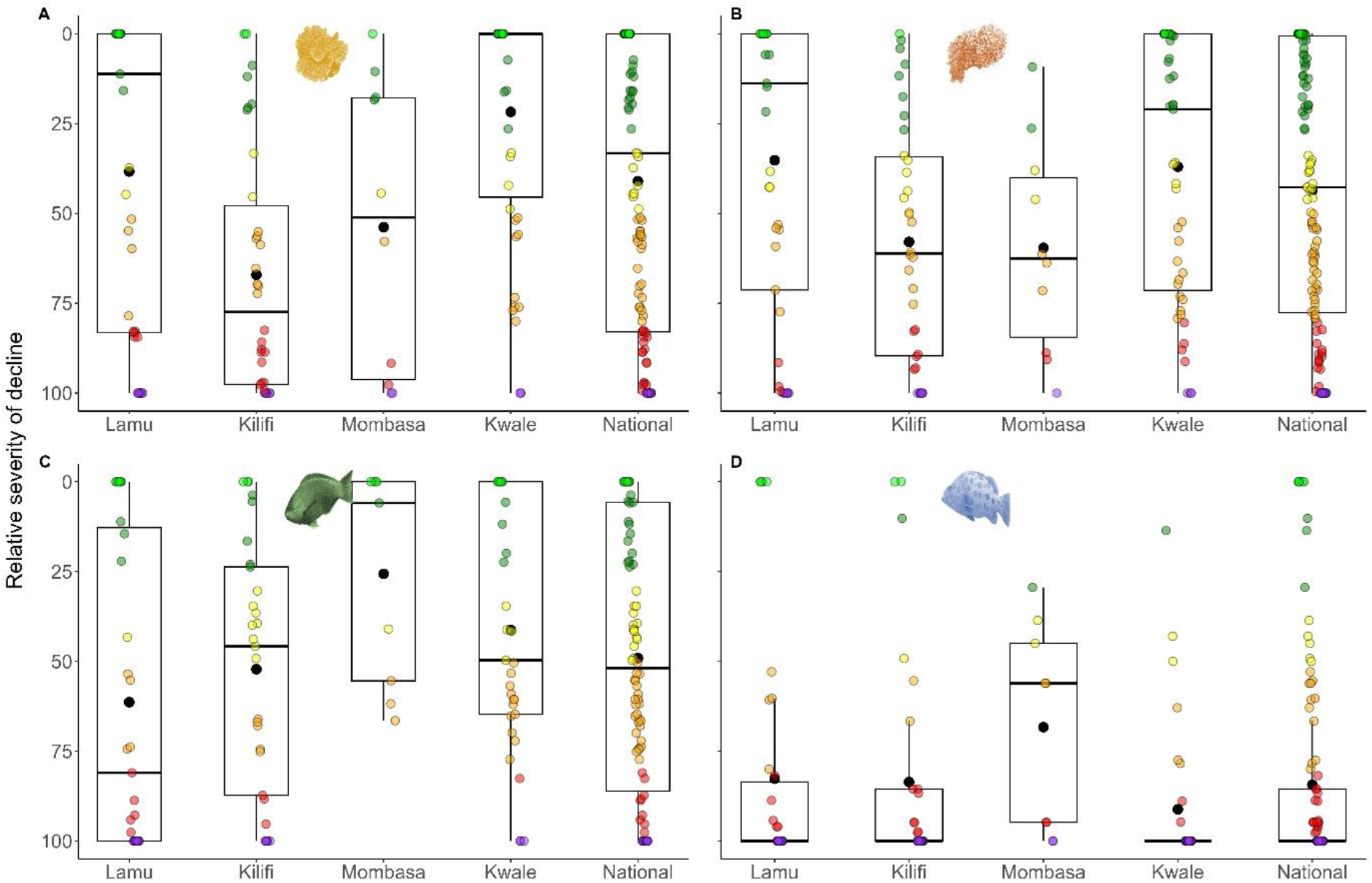
Relative severity of decline over the past 50-years for A) hard coral cover, B) macroalgae-coral ratio, C) parrotfish abundance and D) grouper abundance at coral reef monitoring sites for the entire country and split by the four coastal counties (Lamu, Kilifi, Mombasa, Kwale – north to south) of Kenya. Points are colour coded by relative severity of decline, the horizontal line represents median levels and large, closed, black circle represents the mean in each boxplot. Note, the y-axis is reversed so that values at the top correspond to better condition i.e., low relative severity of decline.

Kwale County had a median relative severity of decline of hard coral over the past 50 years of 0% across its 47 sites, compared to 11% in Lamu (n=41 sites), 51% in Mombasa (n=10) and 77% in Kilifi (n=34) (Fig. 2). For macroalgae-coral ratio, median relative severity of decline varied from 14% in Lamu to 63% in Mombasa (Fig. 2). Parrotfish abundance declined the most in Lamu (median decline=81%, n=31), and the least in Mombasa (median decline = 6%, n=9). For groupers, Mombasa was the only county not to have a 100% median decline (median decline = 56%, n=9).

There were no significant differences in the median relative severity of decline across sites in different management zones for benthic indicators and parrotfish abundance (Hard coral: Kruskal-Wallis (K-W) rank sum test chi-squared = 2.02, p=0.73, macroalgae-coral ratio: K-W chi-squared = 3.7, p=0.44, parrotfish: K-W chi-squared = 6.1, p=0.2) (Fig. 3). However open-access sites had a significantly higher decline in grouper abundance compared to sites with some management (K-W chi-squared = 13.5, p=0.009).

**Figure 3:**
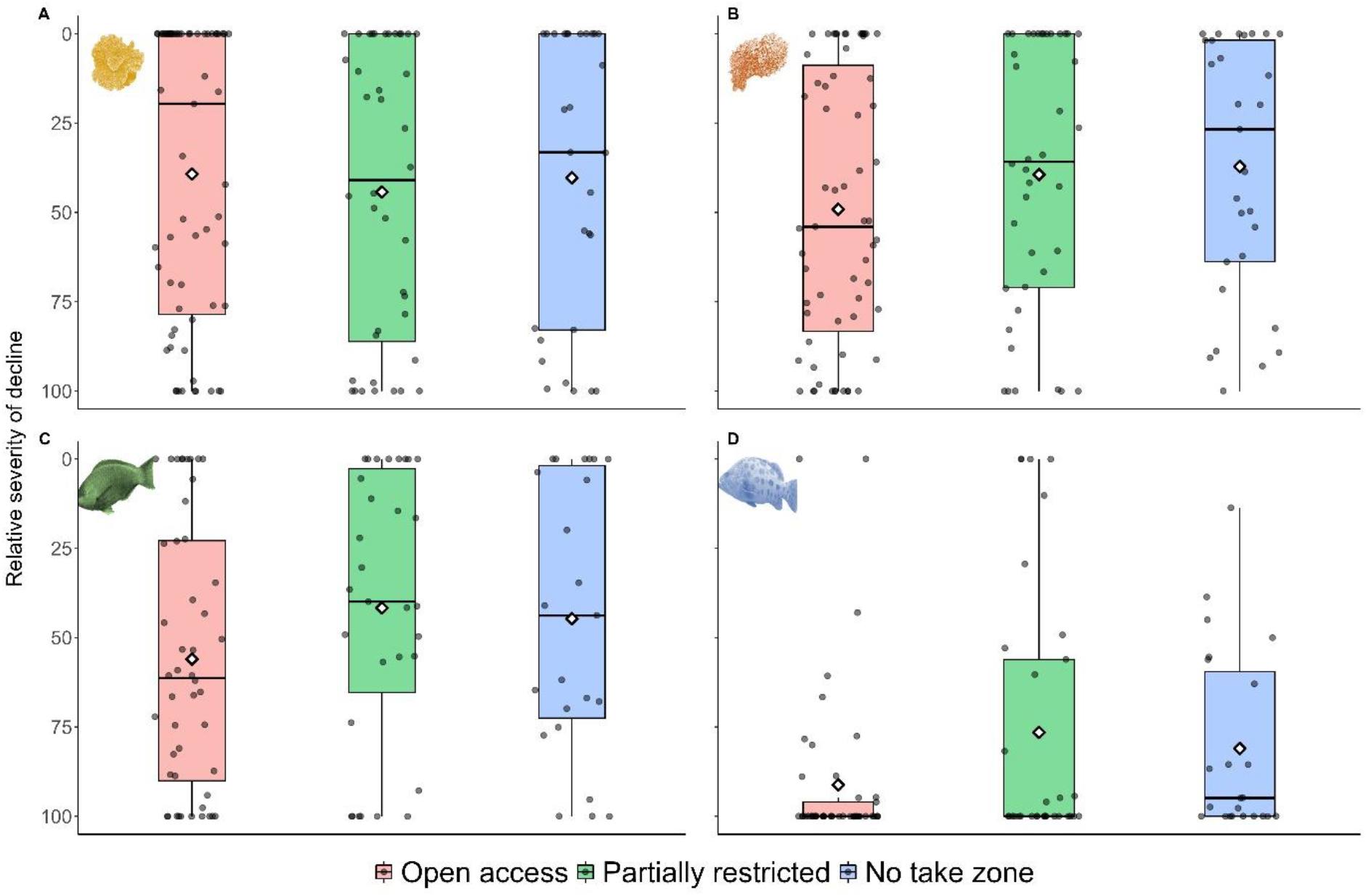
Relative severity of decline over the past 50-years for A) hard coral cover, B) macroalgae-coral ratio, C) parrotfish abundance and D) grouper abundance at coral reef monitoring sites in Kenya, across three different management regimes: no take zone (strict no fishing), partially restricted (artisanal fishing only) and open access (no fishing restrictions). Horizontal line in boxplot represents median levels and large, open diamonds represents the mean. Note, the y-axis is reversed so that values at the top correspond to better condition i.e., decreasing relative severity of decline.

Coral cover and macroalgae-coral ratio in no-take sites in Kisite-Mpunguti (n=9) and Malindi (n=4) MPAs all fared better than corresponding reserve sites (n=6 (Kisite) and n=3 (Malindi)), however the opposite was true for Mombasa Marine Park sites (n=5 in no-take zone, n=4 in reserve) (Fig. S1). Kisite no-take sites also had lower declines for parrotfish and grouper abundance than corresponding reserve sites. Reserve sites in Watamu (n=5) performed better compared to no-take sites (n=5) for both parrotfish and grouper abundance, while in Malindi (n=5 in no-take zone, n=2 in reserve) this was only true for parrotfish.

### Supporting local management decisions

PC1 is strongly correlated with both benthic indicators (hard coral: 0.72, macroalgae-coral ratio:0.67), and PC2 is strongly correlated with both fish abundance indicators (parrotfish: 0.72, grouper: 0.68). The first two principal components explain 69% of the variation in the four indicators of relative severity of decline (Fig. 4). Twenty-eight sites from all four counties were in the upper right ‘healthy’ quadrant, that is, they had above median benthic and fish conditions. Several sites (22) had poor benthic condition but promising fish populations in the lower right quadrant e.g., Dolphin, Drummers and Ch. Mongo Shariff. Twenty-two sites in the upper left quadrant had favourable benthic state but depleted fish populations, such as Shimo la Tewa. Twenty-eight sites were in the bottom-left ‘degraded’ quadrant, due to large declines in both benthic condition and fish abundance.

**Figure 4:**
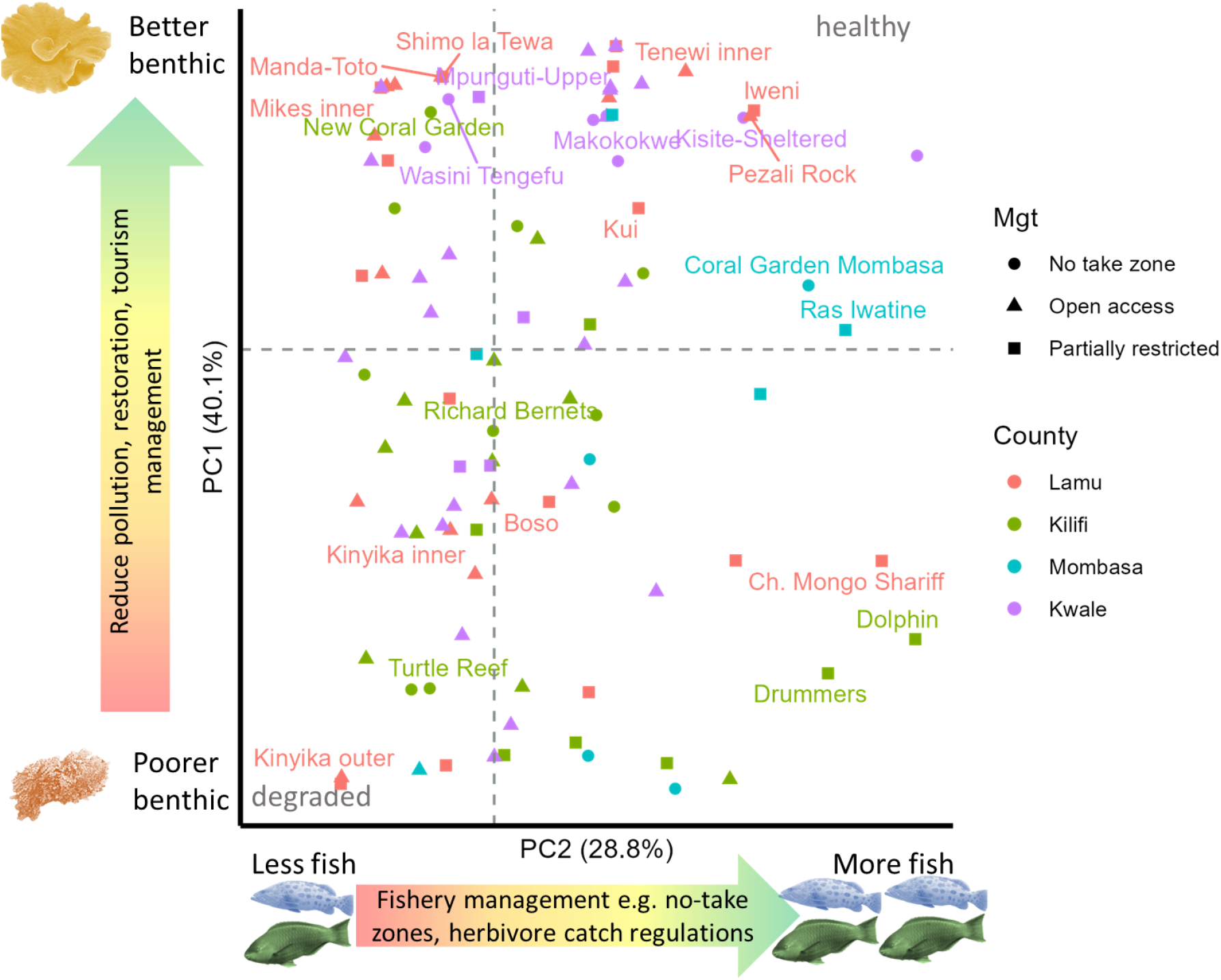
Principal Component Analysis of median relative severity of decline of four indicators (hard coral cover, macroalgae-coral ratio, parrotfish and grouper abundance) over the past 50-years (increasing relative severity of decline in negative directions). PC1 (y-axis) is strongly correlated with both benthic indicators and PC2 (x-axis) is strongly correlated with both fish abundance indicators. Coral reef monitoring sites (n=100) are differentiated by coastal county (colour) and management regime (symbols). No take zone (strict no fishing), open access (no fishing restrictions) and partially restricted (artisanal fishing only). Horizontal and vertical dashed lines depict median PC values, and split space into four quadrants. Management interventions linked to GBF targets are suggested in arrows outside axis. Site name labels shown for select sites with extreme positions within the phase-space, and for sites with hard coral time-series data.

### Trajectory analysis

Out of the 38 sites with suitable time-series data for hard coral cover, 23 sites had a positive median Z-score, indicating that they had above average coral cover for most of their monitoring history. Of these, seven had a score above 1 (substantially above average) (Fig. 5A). Four sites were classified as ‘resistant’ based on our classification criteria (Table 1) (New Coral Garden, Tenewi Inner, Mike’s Inner and Shimo la Tewa) as they maintained positive Z-scores throughout their monitoring history. Seven sites were considered ‘degraded’ because they were consistently below national averages. Nine sites displayed recovery trajectories, while 11 sites displayed declining trajectories. Six sites were ‘resistant’ using macroalgae-coral ratio: Wasini, Kisite East, Upper Mpunguti, New Coral Garden, Old Coral Garden and Mikes Inner (Fig. 5B). A total of nine sites were classified as having undergone a ‘phase-shift’ (coral to algae dominated) using macroalgae-coral ratio e.g., Turtle Reef and Kinyika Inner.

**Table 1:** Reef trajectory classifications using hard coral and macroalgae-coral ratio Z-scores and standard deviation (phase-shifted is based on macroalgae-coral ratio only):

| Classification | Description | Variables and values |
| --- | --- | --- |
| *Resistant* | Always maintained above average condition with minor changes | median Z > 0 & min Z >= ~0 |
| *Degraded* | Consistently below average condition | median Z < 0 & low sd & recent Z direction <= 0.1 |
| *Recovered* | Changed from a below average state at some point to currently having an above average and improving condition | medium/high sd & min Z < ~0 & recent Z direction > -0.1 & recent Z > 0 |
| *Declined* | Changed from an above average condition/state at some point to currently being below average and declining condition | medium/high sd & max Z > ~0 & recent Z direction < 0.1 & recent Z < 0 |
| *Phase-shifted* | using macroalgae-coral ratio only; high variation, starting off moderately above average to currently being below average by a moderate amount. Similar to declined but start at end states must be moderately distant from average. | medium/high sd & recent Z direction < 0.1 & recent Z < -0.5 & first Z > 0.5 |

**Figure 5:**
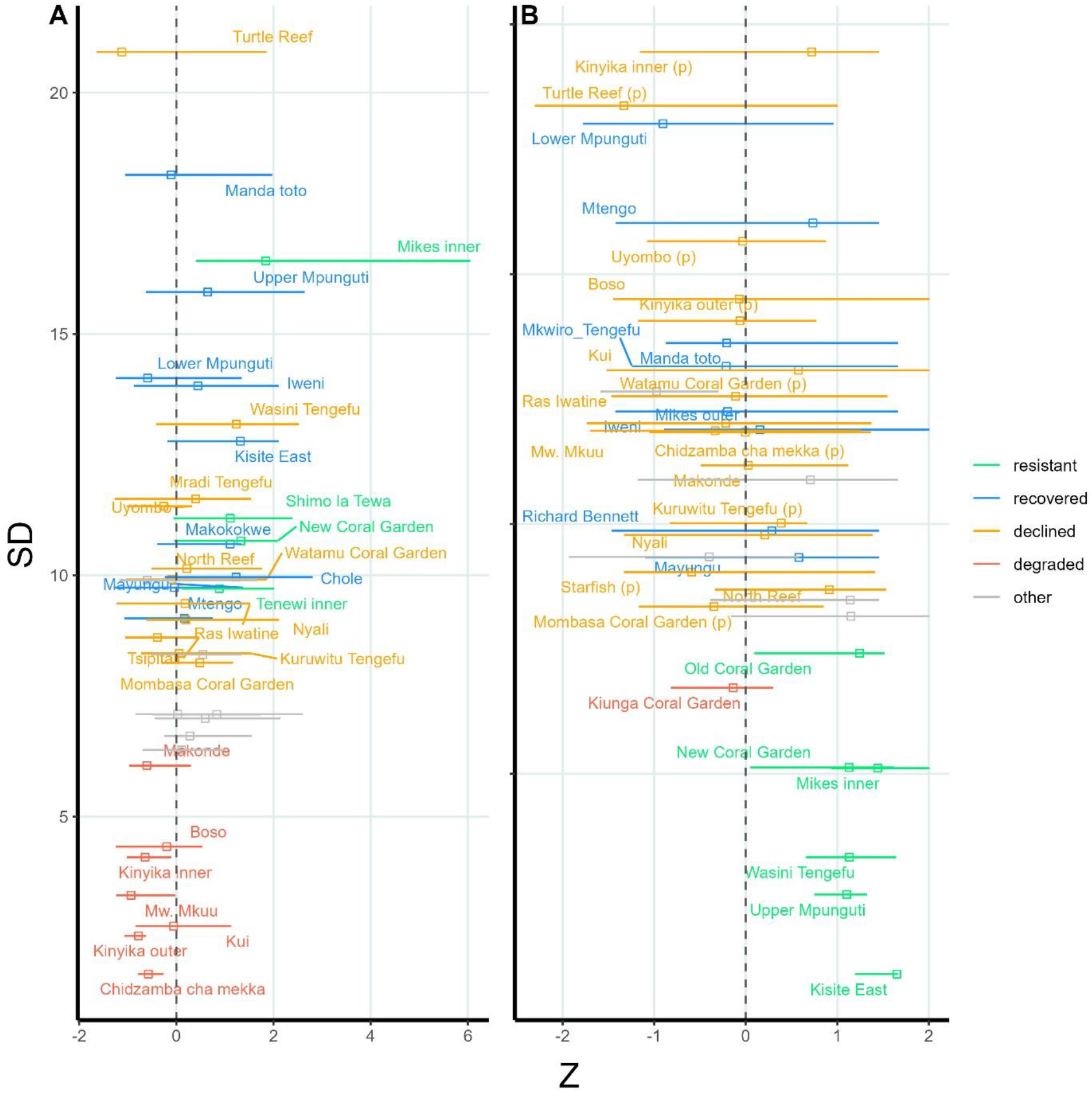
Coral reef monitoring sites in Kenya with long-term time-series data (at least four annual surveys) plotted by the inter-annual variation (standard deviation of the mean of all surveys, y axis) vs the median (open square) and range (horizontal line) of Z-scores across a site’s monitoring history (x axis) for A) hard coral cover (n=38) and B) macroalgae-coral ratio (n=36). Positive Z-scores represent sites which are generally above the national average, while high SD represent sites with large inter-annual variation. Sites are colour coded by trajectory classification (from Table 1). ‘Other’ represents sites with unclassified trajectories. Dashed vertical line indicates Z=0 value. In B, sites annotated as ‘Declined’ with (p) following the site names indicate those which could be considered phase-shifted

## Discussion

### Overview

We show how reef-level data on ecosystem degradation used in the Red List of Ecosystems can support local management decisions for a threatened ecosystem. Our analyses found a general trend of ecological degradation across four key components (hard corals, macroalgae, herbivory, piscivory) of Kenya’s coral reef ecosystem over the past 50 years. This resulted in a Vulnerable risk status under criterion D of the RLE, which is consistent with the result for the broader ecoregion containing northern Tanzania (Obura et al. 2021). At more than half the monitored sites both parrotfish and grouper abundance declined by more than 50%, while coral cover and macroalgae-coral ratio declined by at least 30%. Our results guided recommendations on potential sites to implement management actions which can contribute to meeting relevant GBF targets at a national and global scale, including targets for restoration (Target 2), protection (Target 3), and mitigating and minimising pollution (Target 7).

### Status of Kenya’s coral reefs

The results provide quantitative and standardised estimates of degradation and collapse risk at a reef site scale across a substantial extent of coral reefs in Kenya. Our approach offers a holistic perspective on ecosystem functioning by assessing four critical ecosystem components. There is general trend of degradation, although the degree varied across the system, indicating that site and localised factors have a strong influence on state and recovery (Karisa, 2008; McClanahan et al., 2008; Mwaura et al., 2017; Karisa et al., 2020). Although fish abundance (parrotfish and groupers) showed declines across most sites, a sign of inadequate fisheries regulations and management along the Kenyan coastline (Samoilys et al., 2017), MPAs and LMMAs which restrict fishing and control gears had a significant positive effect on groupers. In Mombasa County, where all sites are protected, declines in both grouper and parrotfish abundance were the lowest. In Kwale County, Kisite MPA has higher fish abundance in no-take zones compared to reserves (where regulated artisanal/traditional fishing is allowed), corroborating studies that demonstrate the effectiveness of no-take zones in Kenya (McClanahan and Mangi, 2000; McClanahan and Arthur, 2001; McClanahan and Graham, 2005; McClanahan et al., 2007; Samoilys et al., 2017). However, the opposite was true for parrotfish in two MPAs (Watamu and Malindi), likely because reserve sites are at deeper locations on the fore reef, compared to the shallow, lagoonal no-take zones.

In theory, control of fishing within MPAs should lead to higher coral cover (Selig and Bruno, 2010; Lamb et al., 2015; Mellin et al., 2016), particularly if new measures curb destructive fishing or recreational practices that damage coral. However, this positive effect of local management was not always detected in our study, likely due to the indiscriminate impact of bleaching events across management zones (Johnson et al., 2022). Kisite-Mpunguti Marine Park and Reserve in Kwale county, which is known for having the most complex reef system and seascape bathymetry, as well as the largest no-take zone in the country (Tuda and Omar, 2012), had sites with the best coral conditions. Sites in the northern MPAs of Watamu and Malindi have experienced more degradation, likely a result of bleaching combined with increased sedimentation and pollution from a major river and coastal development (Samoilys, 1988; van Katwijk et al., 1993; Obura et al., 1994; McClanahan and Obura, 1997; Mwachireya et al., 2015).

The trajectories in coral cover revealed four sites that have some resistance to disturbances i.e., maintain an above average condition over their monitoring history (nine sites if possible sampling inconsistencies are considered). These sites should be a top priority for protection against local threats like fishing and pollution. Sites which showed a recovery trajectory of bouncing back from mortality events may also be a management priority as this characteristic will become increasingly important as climatic conditions continue to change. Managing these sites would help to achieve Target 8 related to building ecosystem resilience to climate change. The seven sites with ‘degraded’ coral cover (low variability and negative median Z-scores) may require management action if important for fishing (e.g., Boso and Kui) or of cultural significance or tourism value (e.g. Kinyika). There was broad consistency between the two benthic indicators. For example, both indicators classified the same six sites as ‘resistant’ after considering uncertainty due to sampling inconsistencies. Macroalgae-coral ratio was also able to provide extra context on the condition of reefs, for example, five sites classified as ‘declined’ and three sites as ‘degraded’ based on hard coral, were characterised as ‘phase-shifted’ using macroalgae-coral ratio (e.g., Kinyika Inner).

### Local management applications for Kenyan coral reefs

A key objective of this study was to develop a method that uses data and indicators from an RLE assessment to inform management at a local spatial scale. Supporting management decisions to tackle local threats such as overfishing, pollution and sedimentation is crucial in Kenya, as the East African coast is projected to have a more favourable climate outlook compared to other regions globally (Beyer et al., 2018; Obura et al., 2021). Using just four indicators, our decision-support framework provided site-level management recommendations related to nine of the 23 GBF targets (Table S3). This site-scale approach mirrors how Kenyan reefs are currently managed, rather than through broader-scale systematic planning (McClanahan et al., 2005; Ban et al., 2011; Maina et al., 2015), and considers the GBF targets that are directly related to the uses and threatening activities in the country. Importantly, sites are spread across most of the phase-space of our framework (Fig 4). We identified several sites from across the country with relatively healthy benthic and fish communities as candidate areas for spatial planning and protection measures to maintain their high biodiversity and ecological integrity, under Targets 1 and 3. Effective fisheries management and enforcement such as no take zones, fishing gear restrictions and species catch restrictions, can contribute to Target 5 to attain sustainable harvesting levels of wild species, and to enable threatened grouper species to recover under Target 4. Sites with poor benthic state but relatively abundant fish populations could be considered as candidates for restoration (e.g. coral gardening) as part of Target 2, particularly as they have herbivorous fish required to keep algae under control, increasing the likelihood of success (Ladd et al., 2018; Ladd and Shantz, 2020; Gouezo et al., 2021; Knoester et al., 2023). Sites with low fish populations and poor benthic condition may benefit from interventions to reduce threats, including tourism management (under Target 9), pollution reduction (Target 7), and/or fishery and herbivore management (Target 5), depending on the context (see Table S3 for full details).

### Guidance for future assessments to support local ecosystem management

Our approach for coral reefs provides several learning outcomes that can be applied by Parties to the CBD to produce site-specific management recommendations as they assess reefs in other regions, as well as other ecosystems. For those countries without an RLE assessment, Reporting on Goal A and Target 1 should involve as a priority, the generation of a national Red List risk status for each ecosystem type (Nicholson et al., 2024). This will include a synthesis of knowledge on the ecosystem types in the country, their distribution, and indicators of change in area and integrity. The analytical approach to achieve this outcome will depend on the data and resources available, but as we’ve done, assessing the input data at the finest resolution possible, provides the flexibility to scale up and down to meet different decision and management needs in the country (Leadley et al., 2022).

Our approach of using indicators on ecosystem state to directly inform action-oriented targets of the GBF can be applied to any ecosystem with spatially explicit ecological data. It requires relating the drivers impacting the ecosystem to the relevant GBF targets. Our management support framework is flexible and allows for nuanced decision-making at high spatial resolutions – to site level where data is robust, and at broader but meaningful spatial-scales where the data is more limited e.g. MPAs or designated use areas. The findings can be incorporated into systematic conservation planning or another decision process (e.g., cost-benefit analysis of potential management actions around LMMAs (Samoilys et al., 2019b)), to be considered alongside other relevant aspects like threats, management costs, feasibility and socio-economic resilience (Joseph et al., 2009; McLeod et al., 2021). For example, reef areas in good condition or with high resistance to ocean warming can be prioritised for conservation based on the exposure and resilience to local threats such as pollution (Target 8).

This comprehensive analysis underscores the value of evaluating multiple, carefully selected biotic indicators for Criterion D of the RLE. While this requires investment in extensive data collection (both spatially and temporally), it is particularly worthwhile when the outcomes are linked to broader applications such as supporting management. Investment in long-term ecosystem monitoring programs that align with the principles of Essential Ocean or Biodiversity Variables (EOV/EBV) have wide-ranging applications beyond the RLE (Miloslavich et al., 2018) e.g. hard coral cover is also used in regional and global status reports and resilience potential assessments (Maynard et al., 2015; Obura et al., 2019; Souter et al., 2021; Obura et al., 2021). Collaborative networks, e.g. the Global Coral Reef Monitoring Network (GCRMN) facilitate the sharing of ecosystem data to build more extensive datasets that offer greater possibilities and stronger outcomes from assessments (Buschke et al., 2023).

## Conclusion

The Global Biodiversity Framework provides a roadmap for how the world can sustain biodiversity and ecosystems. The Red List of Ecosystems has a growing track record of strengthening national environmental policy, legislation, and reporting (Bland et al., 2019; Keith et al., 2023), and has a key role to play in monitoring and meeting the goals and targets of the GBF (Nicholson et al., 2024). Parties to the Convention on Biological Diversity will report on progress for Goal A through the overall risk status of their ecosystems. Our work shows the broader applications the RLE can have across the GBF using the same data, including to support implementation of GBF targets related to ecosystem-based management. Such an approach can be readily applied to other ecosystems where comparable data exists, which should enhance the overall outcomes of the GBF for biodiversity and society.

## Acknowledgments

The authors would like to thank The University of Melbourne and Deakin University for supporting the lead author’s PhD candidature. We would also like to thank Norad (Norwegian Agency for Development Cooperation) for their support towards Mishal’s PhD and CORDIO and its partners’ coral reef monitoring activities through the project ‘Innovating and sharing knowledge for coastal resilience in Eastern Africa’ implemented by CORDIO East Africa. We would also like to thank the Nairobi Convention’s Coral Reef Task Force for Kenya, as well as all the institutions and researchers who contributed their data including Kenya Wildlife Service/Wildlife Research Training Institute, Kenya Marine and Fisheries Research Institute, A Rocha Kenya, CORDIO East Africa and Reefolution, as well as the participants of the various workshops undertaken throughout the National Coral Reef Assessment project.

## Notes

### Competing Interest Statement

The authors have declared no competing interest.

