## Supplementary materials for "Leveraging the Red List of Ecosystems for national action on coral reefs through the Kunming-Montreal Global Biodiversity Framework"

### Methods:

*Relative severity of biotic degradation*

***Table S1:*** Coral reef area in each sub-national zone in Kenya used for calculation of initial values (estimates of 50-years ago) for hard coral cover, macroalgae-coral ratio, parrotfish abundance and grouper abundance.

| **Sub-national zone** | **Coral reef area (km$^2$)** |
|---|---|
| Lamu-Kiunga | 269.2 |
| Watamu-Malindi | 60.6 |
| Mombasa-Kilifi | 54.4 |
| Diani-Chale | 24.5 |
| Msambweni-Kisite | 81.1 |

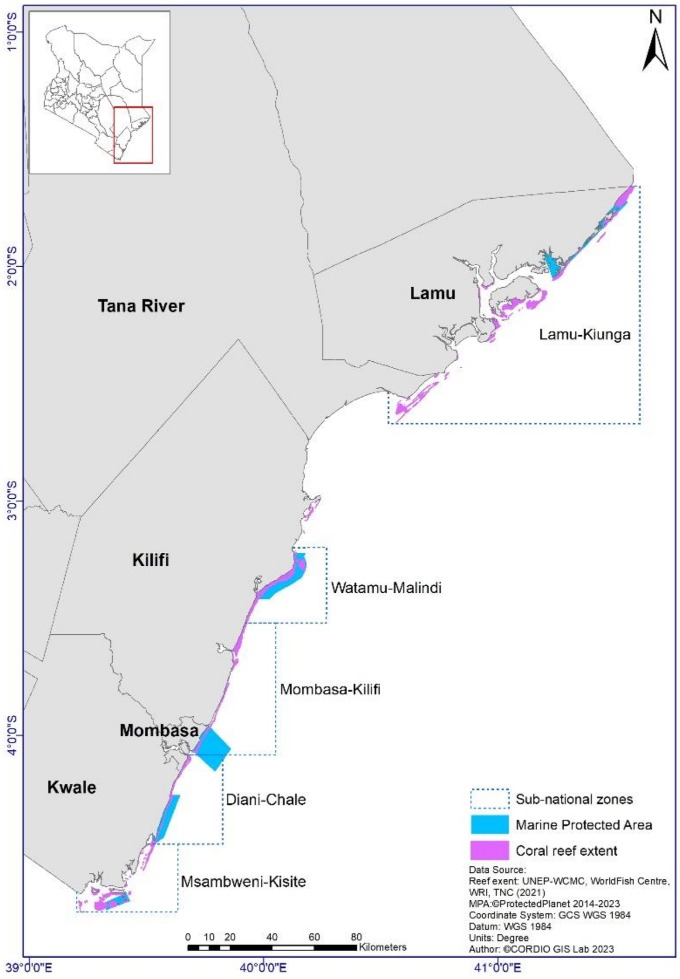


***Figure S1:*** Kenyan coastline showing four coastal counties with coral reefs (Lamu, Kilifi, Mombasa, Kwale) and five sub-national zones used for calculation of initial values (estimates of 50-years ago) for hard coral cover, macroalgae-coral ratio, parrotfish abundance and grouper abundance.

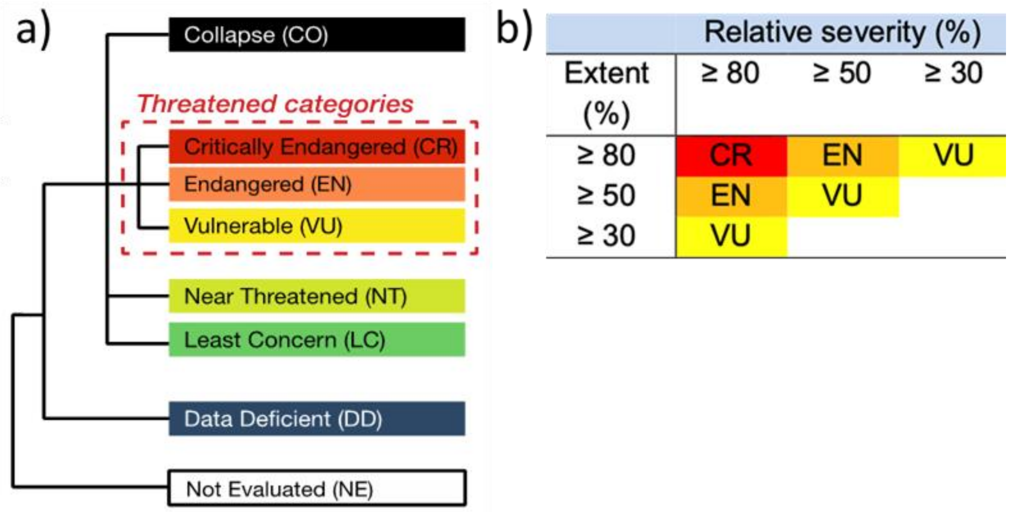


***Figure S2:*** a) Threat categories for ecosystems from the IUCN Red List of Ecosystems (RLE); b) Matrix to determine threat status under Criterion D of the RLE based on ecosystem extent under various relative severity of decline (%) levels. Near Threatened (NT) category was calculated as within 10% of the Vulnerable extent limit for each relative severity level i.e. 27% extent for relative severity ≥ 80, 45% extent for relative severity ≥ 50 and 72% extent for relative severity ≥ 30. Least Concern (LC) was attributed at levels below this.

***Table S2*:** Initial values (mean and intervals/ranges) (estimates of 50-years ago) for hard coral cover, macroalgae-coral ratio, algae-coral ratio, parrotfish abundance and grouper abundance for each sub-national zone. n – number of sites, sd – standard deviation, IQR – inter-quartile range, min – minimum site value, max- maximum site value, cv – coefficient of variation, sd_boot – standard deviation around the mean from bootstrap resampling.

Hard coral:

| zone | n | mean | sd | IQR | min | max | cv | sd_boot |
|---|---|---|---|---|---|---|---|---|
| Lamu-Kiunga | 27 | 15.7 | 5 | 5.6 | 6.3 | 27.5 | 0.32 | 0.94 |
| Mombasa-Kilifi | 14 | 23 | 9.9 | 12.2 | 11.1 | 47.6 | 0.43 | 2.56 |
| Shimoni-Kisite | 5 | 27 | 5.7 | 5.8 | 22.2 | 36.1 | 0.21 | 2.28 |
| Tiwi-Chale | 10 | 13.3 | 7.7 | 11.8 | 4.6 | 25.6 | 0.58 | 2.34 |
| Watamu-Malindi | 18 | 34.4 | 11.1 | 11.7 | 14.3 | 58.9 | 0.32 | 2.57 |

Macroalgae-coral ratio:

| zone | n | mean | sd | IQR | sd_boot |
|---|---|---|---|---|---|
| Lamu-Kiunga | 27 | 0.413 | 0.209 | 0.216 | 0.04 |
| Mombasa-Kilifi | 10 | 0.214 | 0.158 | 0.19 | 0.047 |
| Shimoni-Kisite | 4 | 0.038 | 0.019 | 0.018 | 0.008 |
| Tiwi-Chale | 9 | 0.342 | 0.239 | 0.392 | 0.075 |

| | | | | | |
|---|---|---|---|---|---|
| Watamu-Malindi | 16 | 0.105 | 0.09 | 0.132 | 0.022 |

Fish:

| zone | family | n | mean | sd | IQR | min | max | cv | sd_boot |
|---|---|---|---|---|---|---|---|---|---|
| Kilifi-Mombasa | Grouper | 25 | 166.6 | 204.5 | 80 | 32 | 906.7 | 1.23 | 40.36 |
| Kiunga-Lamu | Grouper | 54 | 114.9 | 171 | 111.2 | 0 | 906.7 | 1.49 | 23.06 |
| Malindi-Watamu | Grouper | 37 | 130.5 | 176.7 | 86.5 | 13 | 906.7 | 1.35 | 28.38 |
| Shimoni-Kisite | Grouper | 24 | 173.7 | 208.1 | 74 | 16.5 | 906.7 | 1.2 | 41.28 |
| Tiwi-Chale | Grouper | 25 | 166.6 | 204.5 | 80 | 32 | 906.7 | 1.23 | 40.36 |
| Kilifi-Mombasa | Parrotfish | 25 | 552.7 | 270.1 | 413.9 | 120 | 1040 | 0.49 | 53.62 |
| Kiunga-Lamu | Parrotfish | 54 | 436.5 | 293.1 | 382.2 | 0 | 1040 | 0.67 | 39.78 |
| Malindi-Watamu | Parrotfish | 37 | 499.8 | 260.2 | 353 | 120 | 1040 | 0.52 | 42.24 |
| Shimoni-Kisite | Parrotfish | 24 | 614 | 261.3 | 429.4 | 120 | 1040 | 0.43 | 52.5 |
| Tiwi-Chale | Parrotfish | 25 | 552.7 | 270.1 | 413.9 | 120 | 1040 | 0.49 | 53.62 |

#### Results:

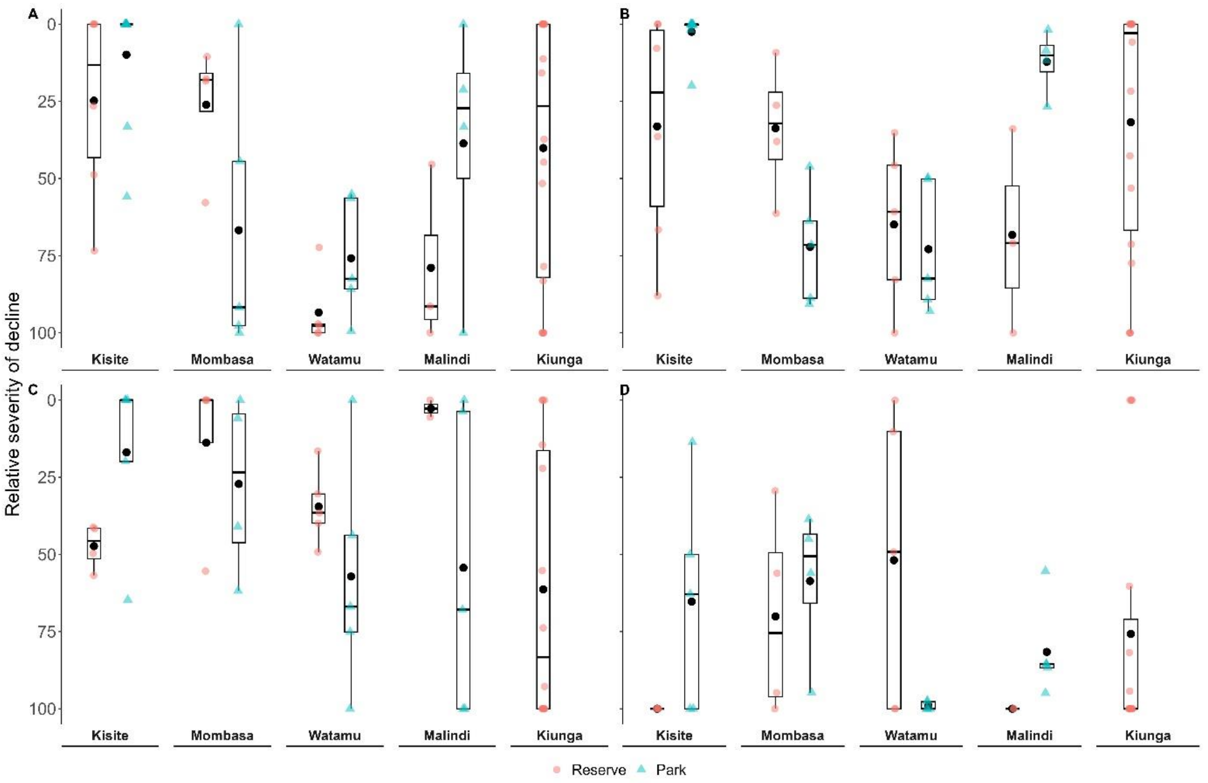


***Figure S3:*** Relative severity of decline over the past 50-years for hard coral cover (A), macroalgae-coral ratio (B), parrotfish abundance (C) and grouper abundance (D) at no take zone (Park) and partially restricted (Reserve) coral reef monitoring sites in five Marine Protected Areas in Kenya. Horizontal line in boxplot represents median levels and large, open diamonds represents the mean. Note, the y-axis is reversed so that values at the top correspond to with better conditions i.e., decreasing relative severity of decline.

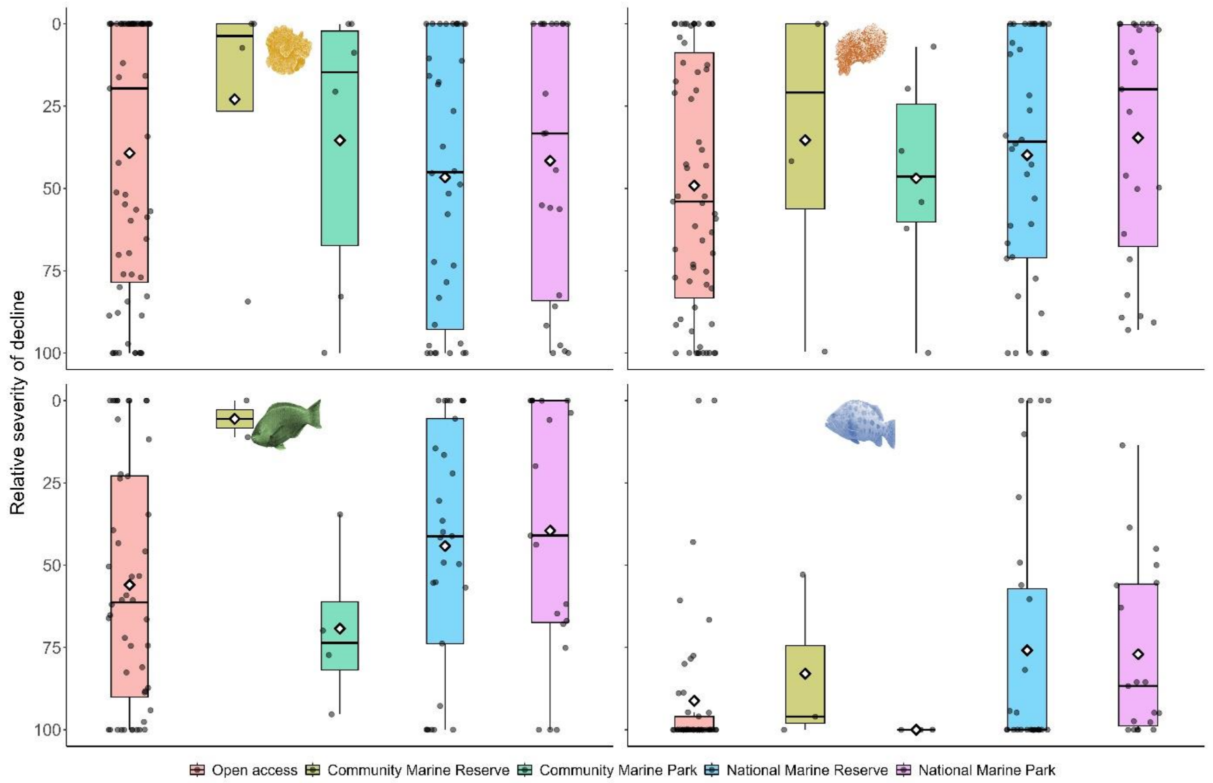


***Figure S4:*** Relative severity of decline over the past 50-years for hard coral cover (top-left), macro algae-coral ratio (top-right), parrotfish abundance (bottom-left) and grouper abundance (bottom-right) at coral reef monitoring sites in Kenya, across five different management regimes: community marine park (strict no fishing), community marine reserve (artisanal fishing only), national marine park (strict no fishing), national marine reserve (artisanal fishing only) and open access (no fishing restrictions). Horizontal line in box-plot represents median levels and large, open diamonds represents the mean. Note, the y-axis is reversed so that values at the top correspond to with better conditions i.e., decreasing relative severity of decline.

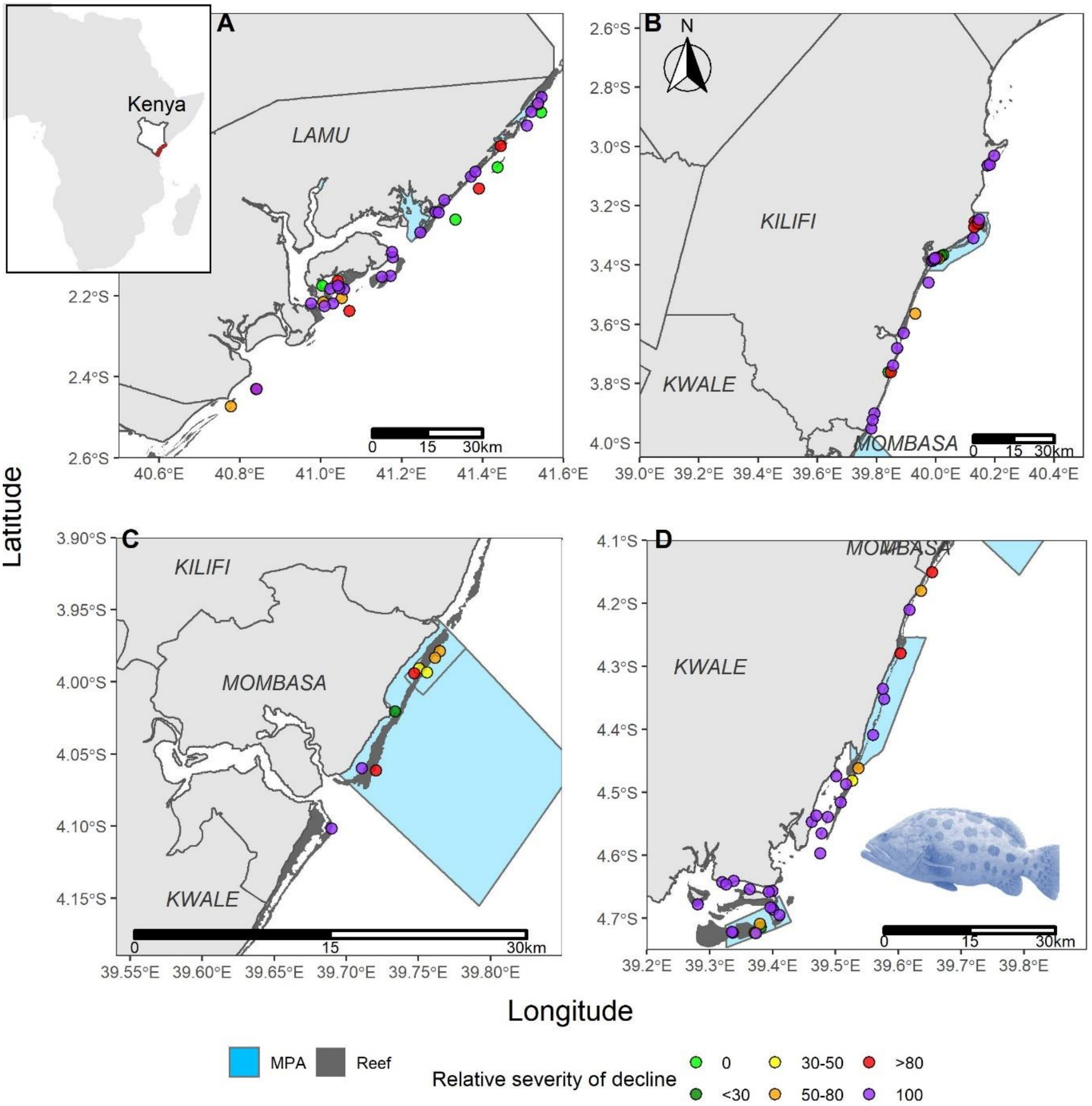


***Figure S5:*** Relative severity of decline classifications over the past 50-years for groupers at coral reef monitoring sites in the four coastal counties of Kenya (A) Lamu, B) Kilifi, C) Mombasa, D) Kwale – north to south). Light blue areas are national Marine Protected Areas (no-take areas and reserves), and dark grey areas represent coral reefs. Sites are coloured based on median relative severity of decline. Inset map on top-left corner shows Kenya with coastal counties highlighted in red.

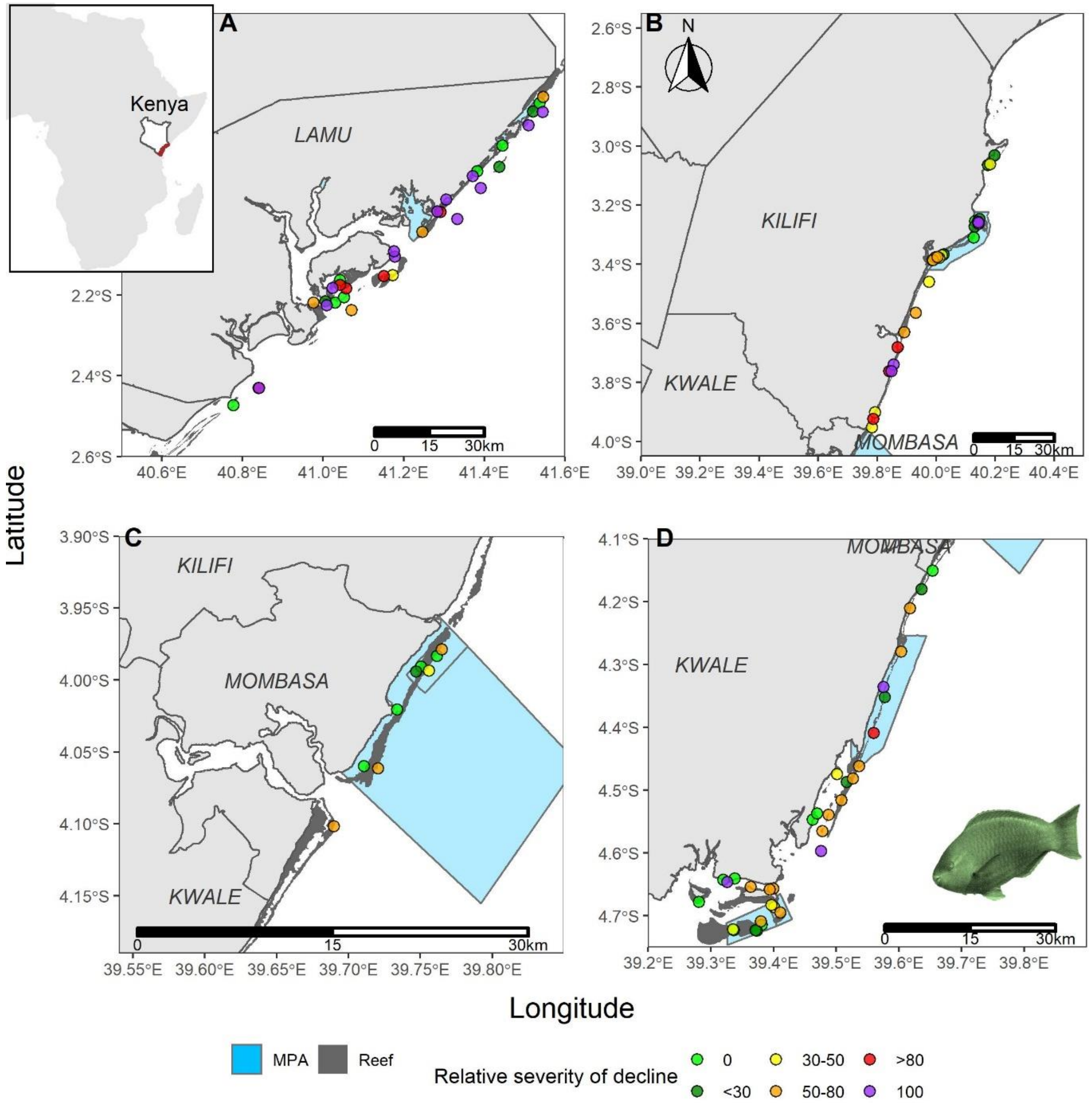


***Figure S6:*** Relative severity of decline classifications over the past 50-years for parrotfish at coral reef monitoring sites in the four coastal counties of Kenya (A) Lamu, B) Kilifi, C) Mombasa, D) Kwale – north to south). Light blue areas are national Marine Protected Areas (no-take areas and reserves), and dark grey areas represent coral reefs. Sites are coloured based on median relative severity of decline. Inset map on top-left corner shows Kenya with coastal counties highlighted in red.

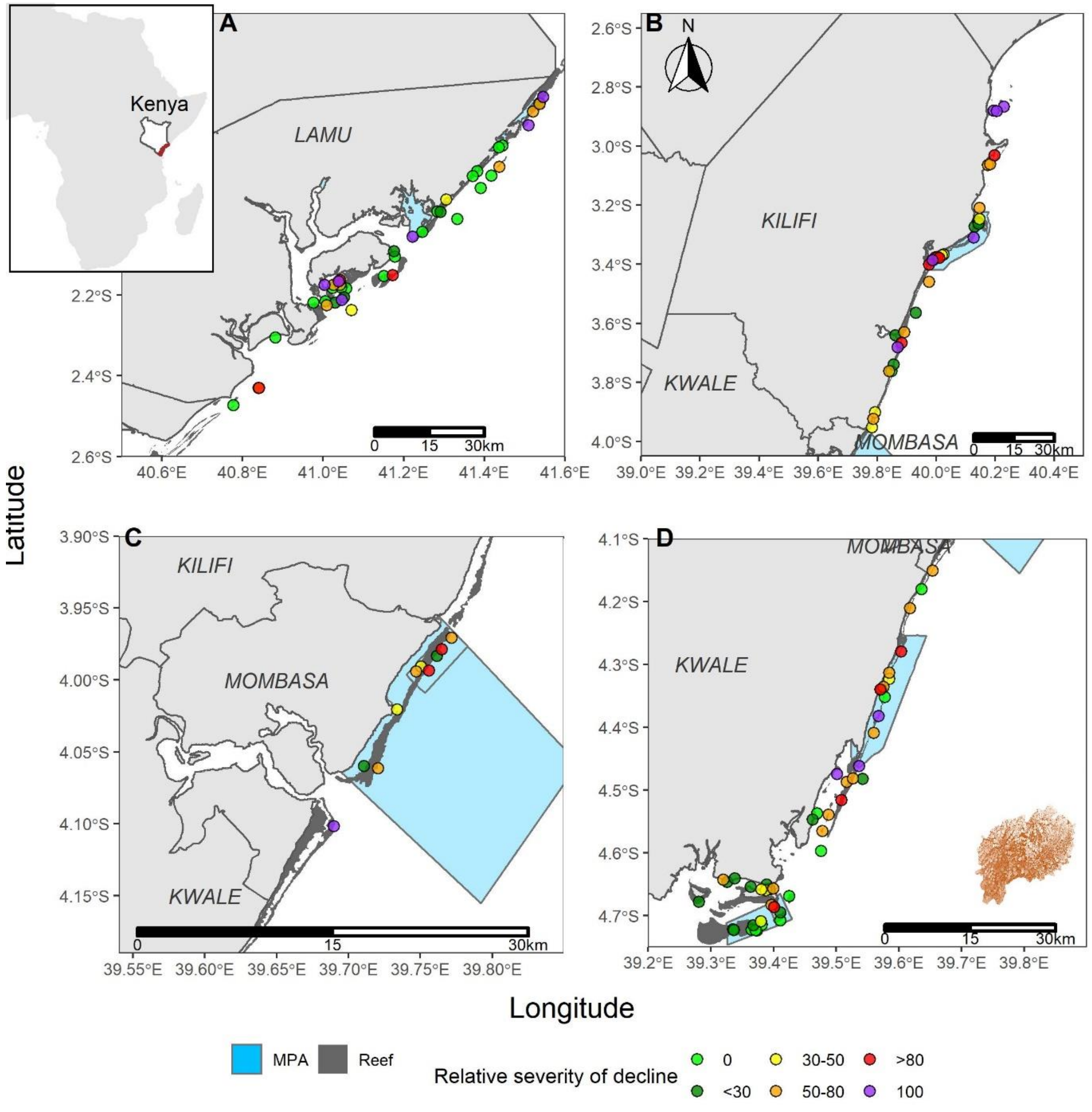


***Figure S7:*** Relative severity of decline classifications over the past 50-years for macroalgae-coral ratio at coral reef monitoring sites in the four coastal counties of Kenya (A) Lamu, B) Kilifi, C) Mombasa, D) Kwale – north to south). Light blue areas are national Marine Protected Areas (no-take areas and reserves), and dark grey areas represent coral reefs. Sites are coloured based on median relative severity of decline. Inset map on top-left corner shows Kenya with coastal counties highlighted in red.

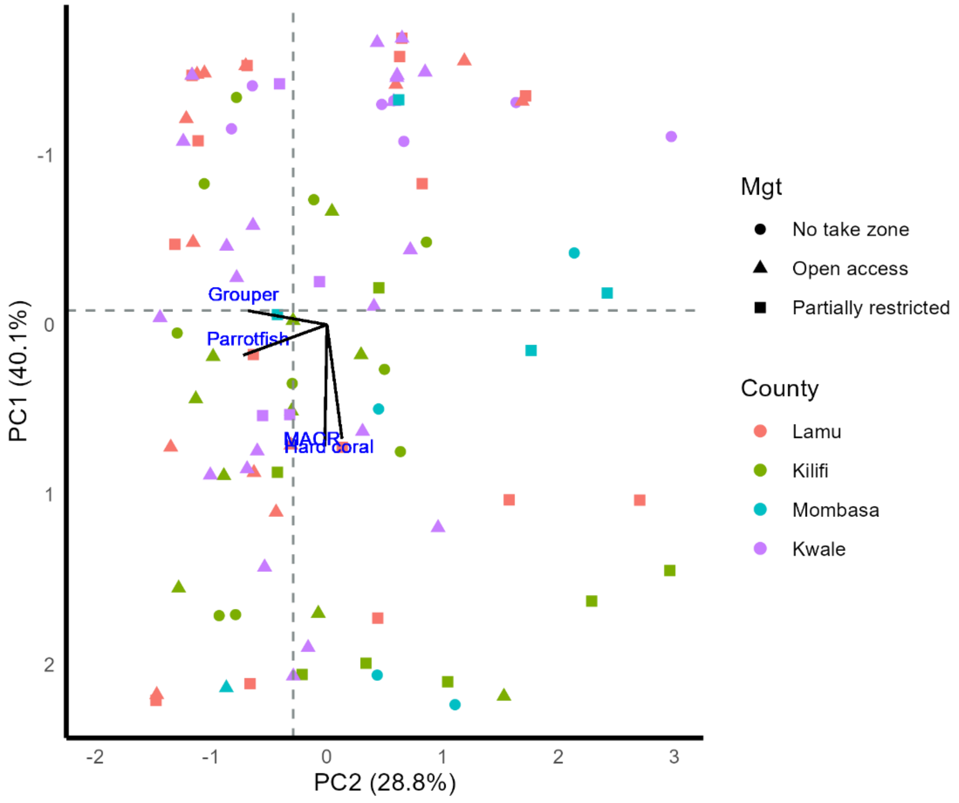


***Figure S8:*** A total of 100 coral reef monitoring sites in Kenya (colour coded by coastal county) arranged using Principal Component Analysis based on median relative severity of decline over the past 50-years for four ecological indicators: hard coral cover, macroalgae-coral ratio, parrotfish and grouper abundance. Shapes based on three different management regimes: no take zone (strict no fishing), open access (no fishing restrictions) and partially restricted (artisanal fishing only). PC1 is strongly correlated with both benthic indicators (HC:0.71, MACR:0.65), and PC2 (Parrot: 0.71, Grouper: 0.69) is strongly correlated with both fish indicators, with increasing relative severity of decline in the positive directions. Horizontal and vertical dashed lines depict median PC values, and split space into four quadrants.

***Table S3*:** Linking practical management interventions for coral reefs to relevant targets of the Global Biodiversity Framework. We explain how the ecological indicators used in this study can be considered through the management support framework to inform implementation of the targets.

| **Management intervention** | **Target** | **Target summary** | **Indicator(s)** | **Link between framework and target/ sites to prioritise** |
|---|---|---|---|---|
| Conservation | Target 1– Spatial planning | All areas under spatial planning or management processes to reduce loss of important areas to zero | All | Sites in a healthy state (good fish and benthic state) are ''areas of high ecological integrity'' or ''important biodiversity areas'', particularly those with resistant or recovery qualities. |
| Restoration (artificial) | Target 2– Restoration | At least 30% of areas ecosystems are under effective restoration | Coral cover, Macroalgae-coral ratio and herbivore abundance | Focus on sites which have low coral cover, but with moderate to high herbivore abundance. Ideally consider those which also have low macroalgae cover. Could target sites which have phase-shifted or declined to attempt to reverse the trend. Could also prioritise |

| | | | | |
|---|---|---|---|---|
| | | | | sites of cultural significance or tourism value. |
| Marine Protected Areas and other effective area-based conservation measures | Target 3–Protection | At least 30% of marine and coastal areas are conserved and managed through protected areas | All | Identify sites outside of existing protected area networks that are in a 'healthy' state for both fish and benthic indicators (as areas of importance for biodiversity and ecosystem functions), particularly those with resistant or recovery qualities. |
| Fisheries management e.g. gear restrictions (mesh sizes and hook sizes) and species restrictions | Target 4–Species<br><br>Target 5–Wild species use | Halt human induced extinction of known threatened species and for the recovery and conservation of species<br><br>Ensure that the use, harvesting and trade of wild species is sustainable, safe and legal | Grouper abundance<br><br>Coral cover; fish abundance | Manage depleted sites with low grouper populations for recovery, and conserve or sustainably manage fishing at sites with existing high populations, particularly those outside of existing fishery management. Additional considerations include targeting deeper and more structurally complex sites (or sites with high coral cover). |
| Pollution management | Target 7–Pollution | Reduce nutrient, chemical and plastic pollution and its impact to levels that are not harmful to biodiversity and ecosystem functions and services | Macroalgae-coral ratio | Actively protect sites with low macroalgae-coral ratio to maintain this state by ensuring nutrient pollution levels are managed/remain low. High macroalgae-coral ratio (particularly phase-shifted) sites should be investigated to identify contributing threats, including pollution. |
| Herbivore management | Target 8–Climate change | Minimize the impact of climate change and ocean acidification on biodiversity and increase its resilience | Parrotfish abundance, coral cover, macroalgae-coral ratio | Prioritise sites with high herbivore abundance, high coral cover, and low macroalgae-coral ratio. Also, sites which have proven resilience qualities - resistant sites or sites which have shown a recovery trajectory. |

| | | | | |
|---|---|---|---|---|
| Tourism management - no-anchoring, diving/snorkelling related damage | Target 9– Benefits | Ensure that the management and use of wild species are sustainable, and provide social, economic and environmental benefits | Coral cover | Sites with high coral cover can be maintained or improved by managing damaging/destructive practices, particularly anchoring and diving. Sites with high tourism visitation that have declined should also be actively managed, |

***Table S4***: National criterion D results for each indicator – proportion of sites (extent) across Kenya in each relative severity classes (classes range from 0 to <30, and from 30, 50 or 80 to 100). n – number of sites, MACR – macroalgae-coral ratio.

| relative severity (%) | **<30** | | **≥30** | | **≥50** | | **≥80** | |
|---|---|---|---|---|---|---|---|---|
| | extent | n | extent | n | extent | n | extent | n |
| Hard coral | 49.2 | 65 | 50.8 | 67 | 43.9 | 58 | 28.0 | 37 |
| MACR | 43.2 | 57 | 56.8 | 75 | 45.5 | 60 | 23.5 | 31 |
| Parrotfish | 35.3 | 36 | 64.7 | 66 | 51.0 | 52 | 27.5 | 28 |
| Grouper | 8.6 | 9 | 91.4 | 96 | 87.6 | 92 | 77.1 | 81 |

***Table S5***: County level RLE Criterion D results for each indicator – proportion of sites (extent) across Kenyan counties in each relative severity classes (classes range from 0 to <30, and from 30, 50 or 80 to 100). n – number of sites, MACR – macroalgae-coral ratio.

| | | **Lamu** | | **Kilifi** | | **Mombasa** | | **Kwale** | |
|---|---|---|---|---|---|---|---|---|---|
| Indicator | Rel severity of decline | extent | n | extent | n | extent | n | extent | n |
| Hard Coral | <30 | 53.7 | 22 | 20.6 | 7 | 40.0 | 4 | 68.1 | 32 |
| | ≥30 | 46.3 | 19 | 79.4 | 27 | 60.0 | 6 | 31.9 | 15 |
| | ≥50 | 41.5 | 17 | 73.5 | 25 | 50.0 | 5 | 23.4 | 11 |
| | ≥80 | 31.7 | 13 | 50.0 | 17 | 40.0 | 4 | 6.4 | 3 |
| MACR | <30 | 56.1 | 23 | 23.5 | 8 | 20.0 | 2 | 51.1 | 24 |
| | ≥30 | 43.9 | 18 | 76.5 | 26 | 80.0 | 8 | 48.9 | 23 |
| | ≥50 | 36.6 | 15 | 58.8 | 20 | 60.0 | 6 | 40.4 | 19 |
| | ≥80 | 22.0 | 9 | 35.3 | 12 | 30.0 | 3 | 14.9 | 7 |
| Parrotfish | <30 | 32.3 | 10 | 27.6 | 8 | 55.6 | 5 | 39.4 | 13 |
| | ≥30 | 67.7 | 21 | 72.4 | 21 | 44.4 | 4 | 60.6 | 20 |
| | ≥50 | 64.5 | 20 | 44.8 | 13 | 33.3 | 3 | 48.5 | 16 |
| | ≥80 | 51.6 | 16 | 27.6 | 8 | 0.0 | 0 | 12.1 | 4 |
| Grouper | <30 | 11.8 | 4 | 10.3 | 3 | 11.1 | 1 | 3.0 | 1 |
| | ≥30 | 88.2 | 30 | 89.7 | 26 | 88.9 | 8 | 97.0 | 32 |
| | ≥50 | 88.2 | 30 | 86.2 | 25 | 66.7 | 6 | 93.9 | 31 |
| | ≥80 | 79.4 | 27 | 79.3 | 23 | 44.4 | 4 | 81.8 | 27 |

Temporal trend analysis

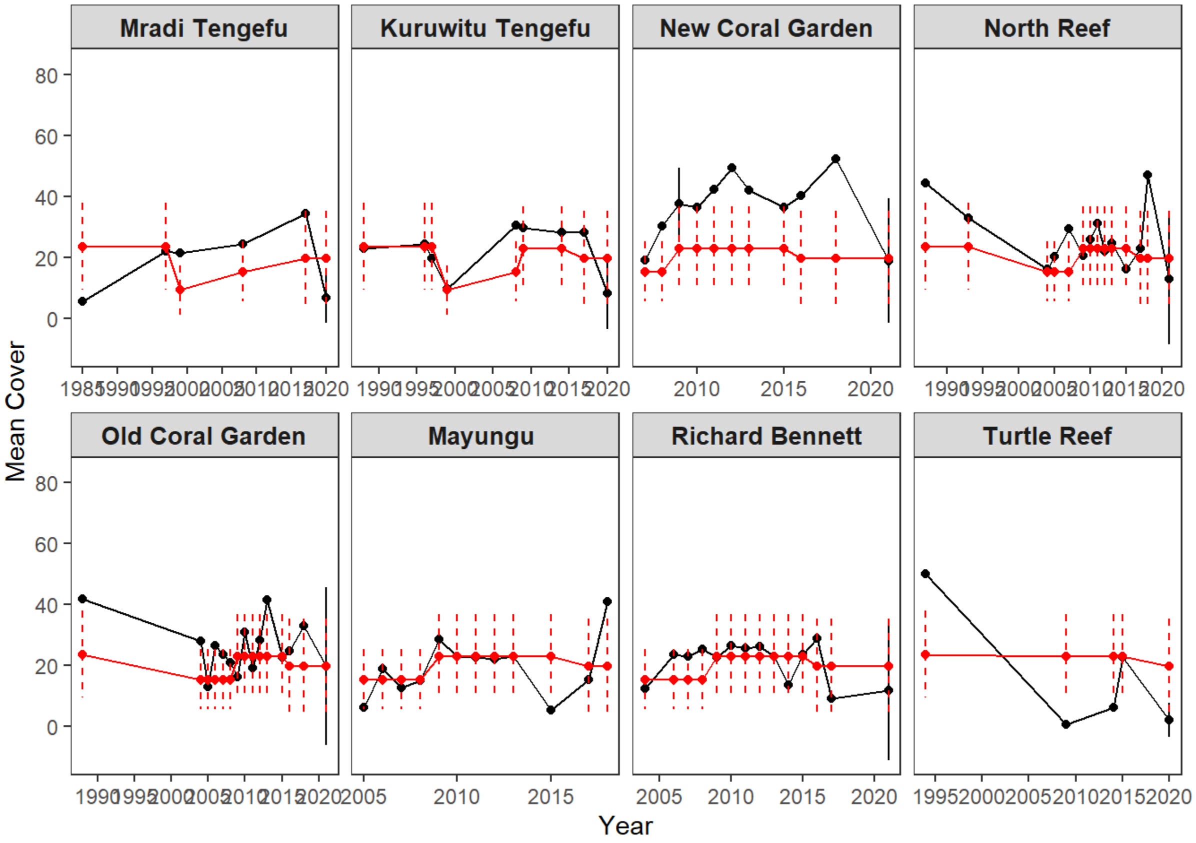


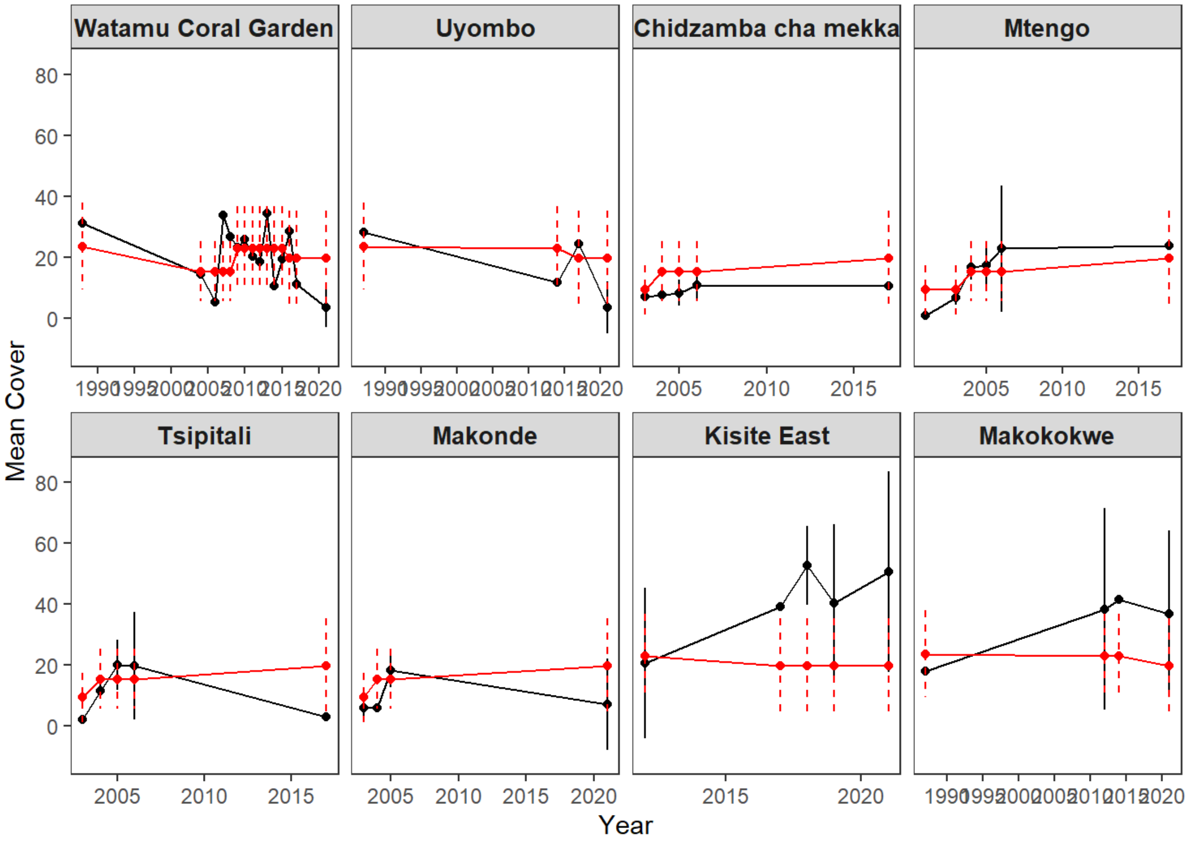

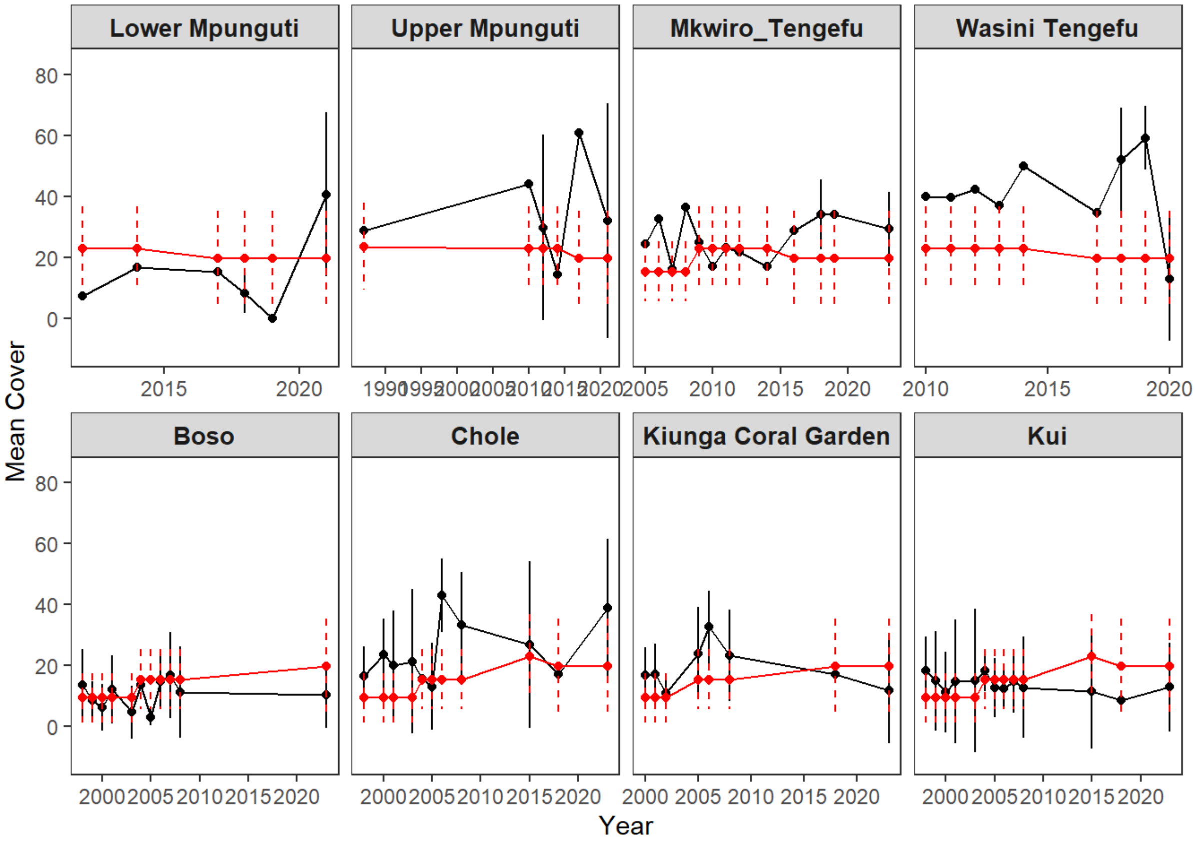
Lower Mpunguti
Upper Mpunguti
Mkwiro_Tengefu
Wasini Tengefu
Boso
Chole
Kiunga Coral Garden
Kui
Mean Cover
Year

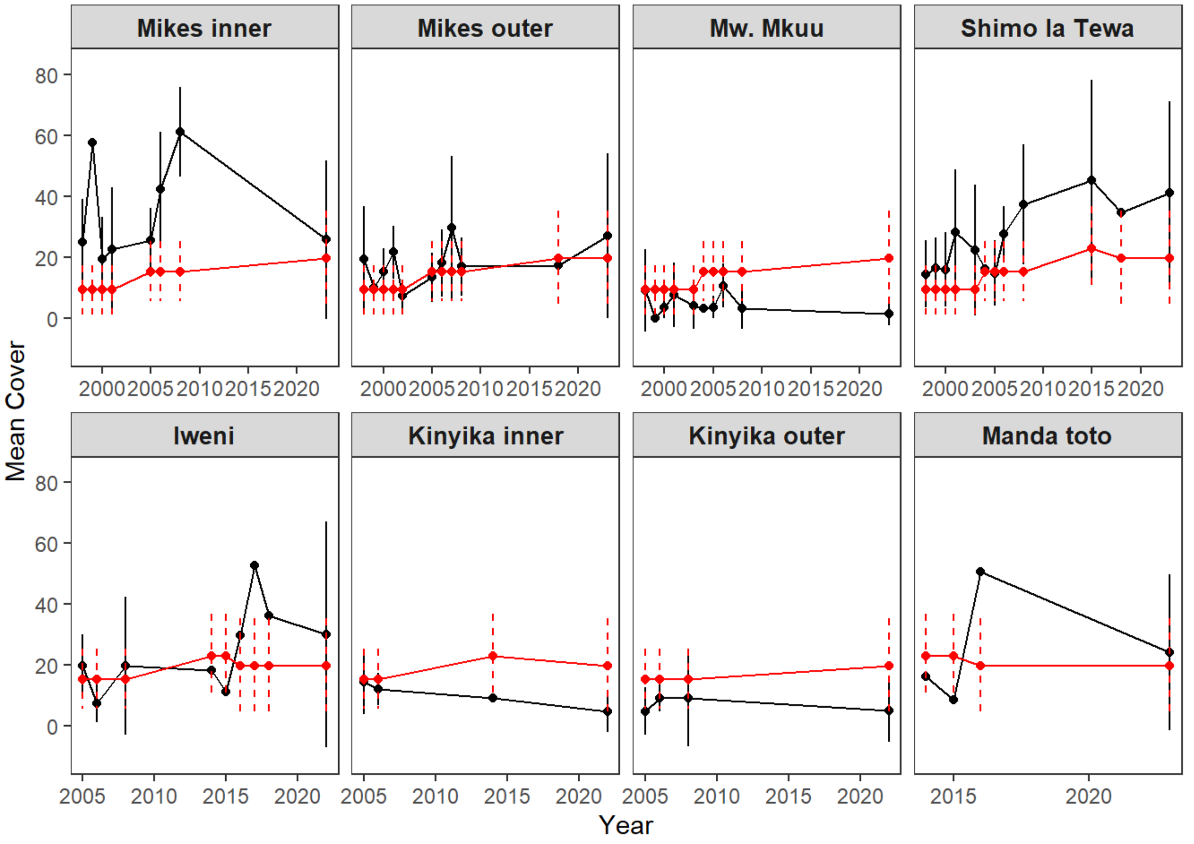
Mikes inner
Mikes outer
Mw. Mkuu
Shimo la Tewa
Iweni
Kinyika inner
Kinyika outer
Manda toto
Mean Cover
Year

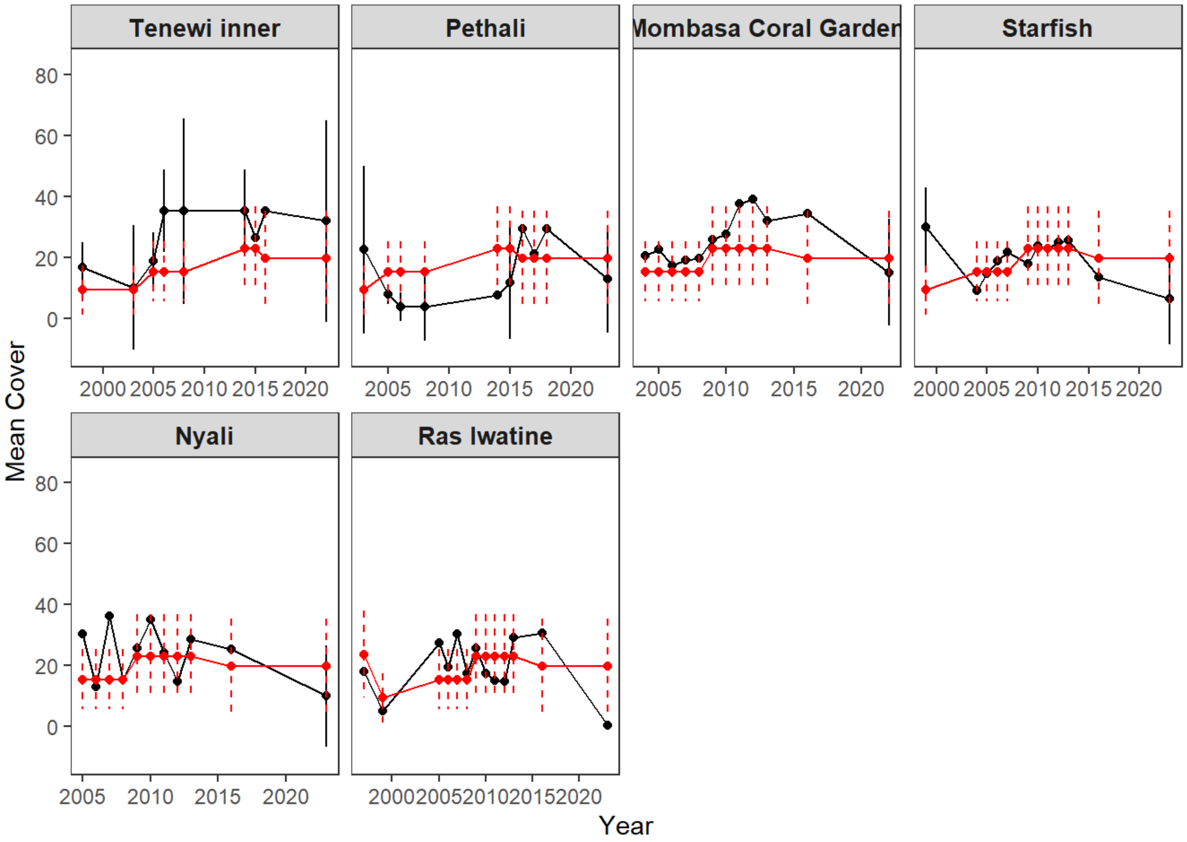


**Figure S14**: Trends in mean hard coral cover (%) at coral reef monitoring sites in Kenya with long-term time-series data (at least four annual surveys, n=38). Black lines and points represent site means and standard deviations, and red lines represent national averages with dashed red vertical lines representing standard deviations (across all sites monitored within that bin).

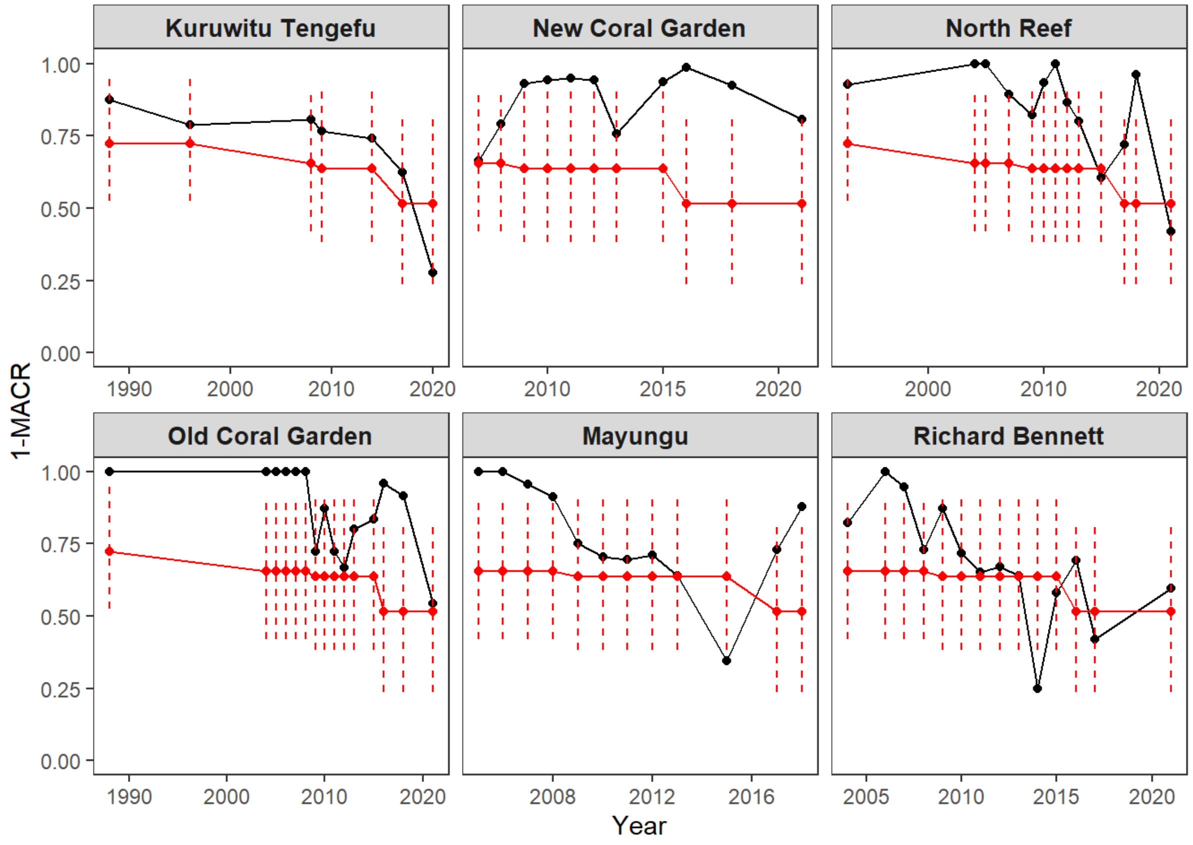
Kuruwitu Tengefu
New Coral Garden
North Reef
Old Coral Garden
Mayungu
Richard Bennett
1-MACR
Year

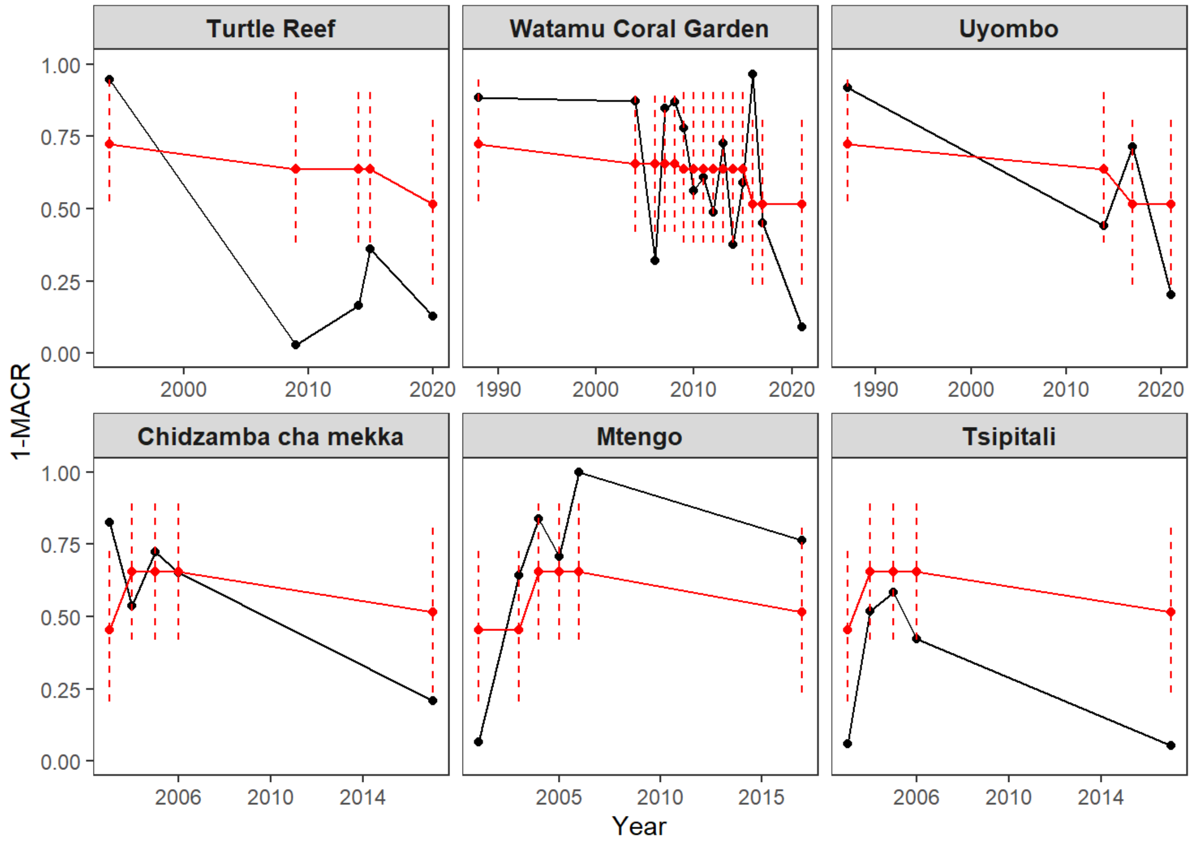
Turtle Reef
Watamu Coral Garden
Uyombo
Chidzamba cha mekka
Mtengo
Tsipitali
1-MACR
Year

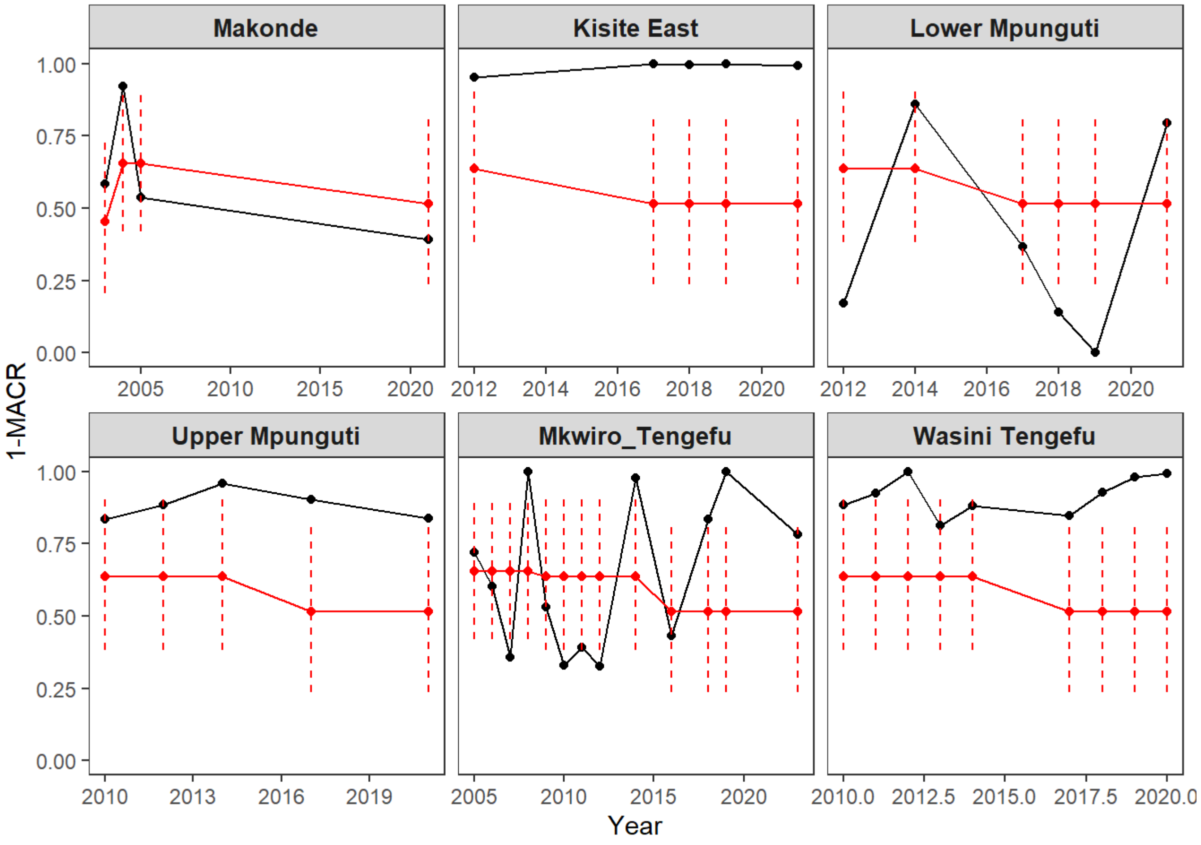
Makonde
Kisite East
Lower Mpunguti
Upper Mpunguti
Mkwiro_Tengefu
Wasini Tengefu
1-MACR
Year

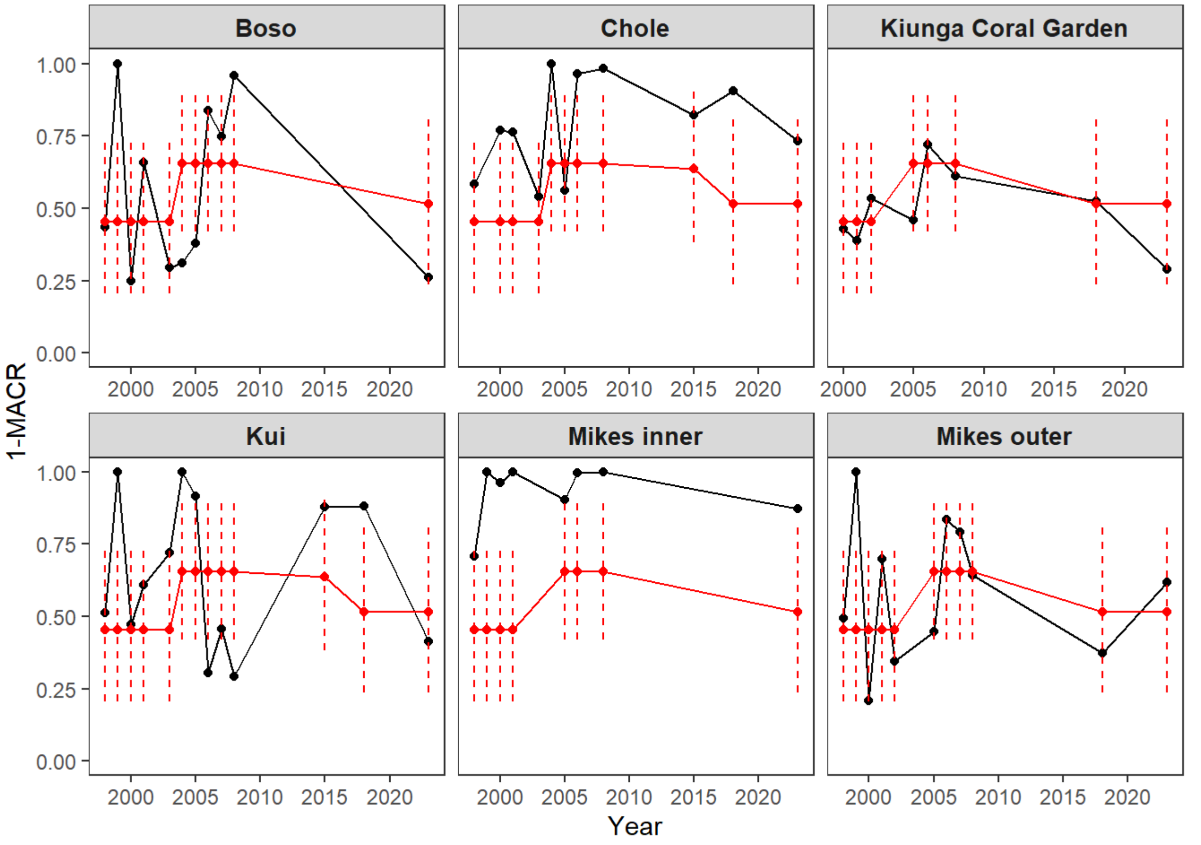
Boso
Chole
Kiunga Coral Garden
Kui
Mikes inner
Mikes outer
1-MACR
Year

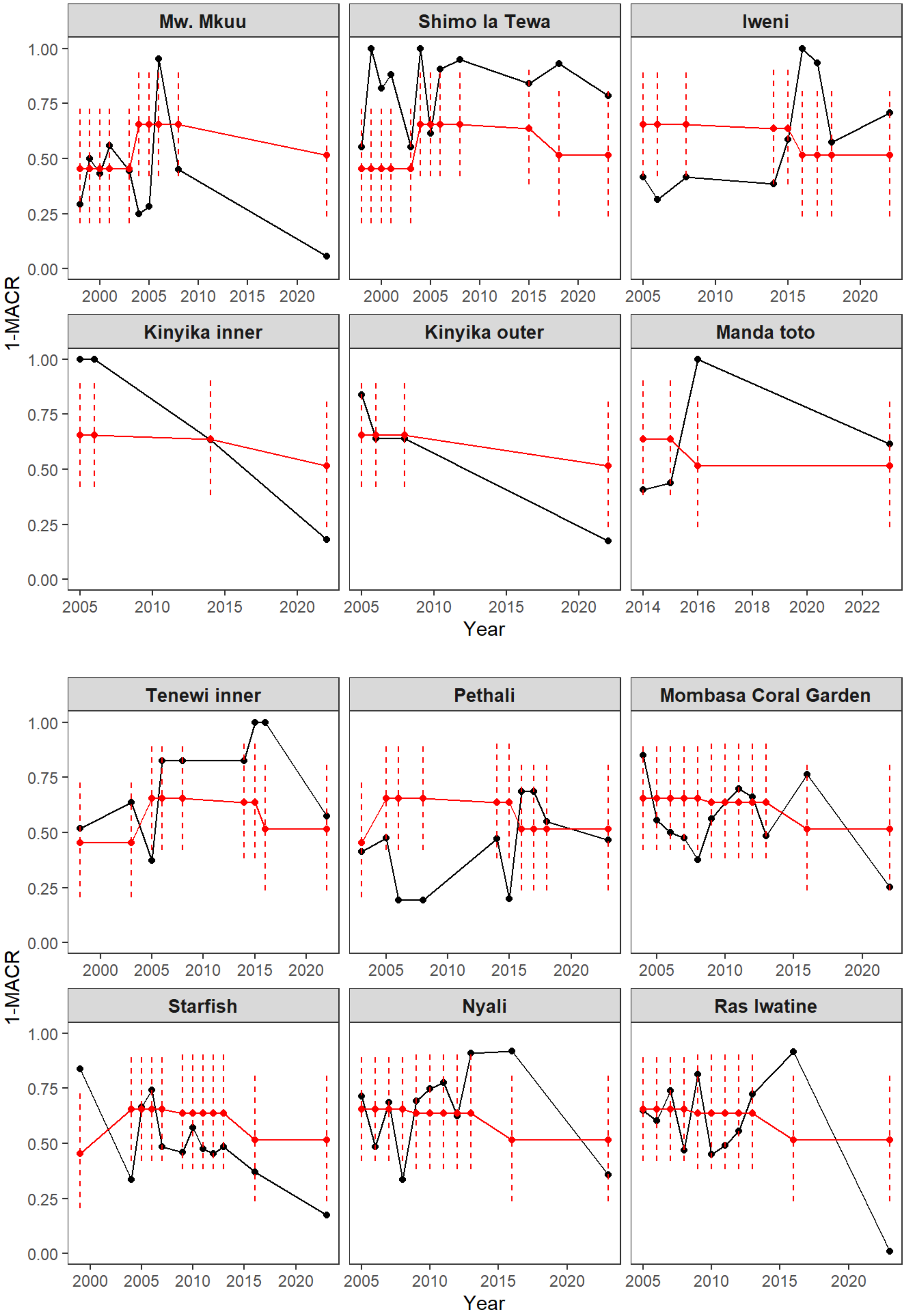


**Figure S15**: Trends in mean macroalgae coral cover ratio (MACR) at coral reef monitoring sites in Kenya with long-term time-series data (at least four annual surveys, n=36). Black lines and points represent 1-site means and standard deviations, and red lines represent national averages with dashed red vertical lines representing standard deviations (across all sites monitored within that bin).
